# Chemically induced cell plasticity enables the generation of high-fidelity embryo model

**DOI:** 10.1101/2024.06.20.598030

**Authors:** Huanhuan Li, Jiahui Huang, Wei Guan, Jinyi Wu, Haiping Luo, Litao Chang, Haiyong Zhao, Chuanxin Chen, Yake Gao, Jian Zhang, José C. R. Silva

## Abstract

Embryo models open exciting opportunities for improving our understanding of development and advancing medicine. However, current models depend on intricate procedures and the integration of separately generated cell types. Here, we aimed to establish a chemical-only strategy to induce cellular plasticity and generate embryo founder-like cells. Within 60 hours, chemical-induced pluripotent stem cells activated early lineage specifiers, *Cdx2* and *Gata6*, while maintaining *Oct4/ Pou5f1* expression. Following a second 36-hour chemical treatment, founder-like cells developed into all blastocyst fates and self-assembled into embryo-like structures. These progressed into high-fidelity late-streak gastrulation stages within 4.5 days, displaying primitive streak formation via epithelial-to-mesenchymal transition, along with germ-layer tissues, amnion, and ectoplacental cone. Fate trajectories and molecular signatures from founder-like cells to chemically-induced embryo model (ci-MO) were defined, revealing remarkable similarity up to Embryonic day (E) 7.5 embryo. Summarily, ci-MO offers a straightforward, rapid, small-molecule-only, and high-fidelity approach to study embryonic development and cell fate specification.

**Highlights:**

- Generation of a direct, efficient, and sophisticated chemically induced embryo model (ci-MO)
- Two-pulse chemical induction reprograms ESCs into founder-like cells capable of developing all embryo fates
- Gastrulating ci-MOs display high morphological and molecular accuracy
- ci-MOs reconstruct embryo developmental trajectories

## Introduction

*In vitro,* cell types akin to the three blastocyst fates have demonstrated the ability to self-organize into early embryo-like structures (Harrison et al., 2017; Rivron et al., 2018). This discovery, along with the realization that these structures, and embryonic stem cells (ESC) alone, can progress into gastrula-like embryo models and gastruloids respectively, mimicking aspects of development such as body plan, marked a significant advancement towards recreating embryo development in a petri dish (Amadei et al., 2021; Beccari et al., 2018). Current state-of-the-art methods involve mixing ESCs with either representative stem cell lines of extra-embryonic endoderm (XEN) /primitive endoderm (PrE)-like and trophoblast stem cells (TSC), or with cells derived from ESC induction using specific culture conditions or defined transgenes (Amadei et al., 2022; Lau et al., 2022; Tarazi et al., 2022). The latter uses the exogenous expression of *Gata6*/ *Gata4* or *Cdx2* to make PrE-like or TE-like cells respectively. The transcriptional trajectory of ESCs induced with *Gata6* shows that ESCs transdifferentiate directly into a PrE-like fate (Li et al., 2023).

Another approach to generate embryo models could be via using cells with higher cell potency, capable of producing all three blastocyst fates. Reprogramming to a developmental stage earlier than that of induced pluripotent stem cells (iPSCs) has been suggested by increasing *Stat3* activation (Stuart et al., 2019), and this has been demonstrated with the generation of morula-like cells (MLC) from ESCs (Li et al., 2023). Interestingly, observed co-expression of the seemingly antagonistic transcription factors *Oct4/Nanog* and *Gata6* was associated with an earlier developmental stage and greater plasticity compared to pluripotent stem cells (Guo et al., 2024; Li et al., 2023; Linneberg-Agerholm et al., 2023; Perera et al., 2022; Redo-Riveiro et al., 2024; Schrode et al., 2014).

Small molecule compounds offer advantages for the directional manipulation of cell identity, presenting great potential for regenerative medicine. These small molecules enable precise control of cell fate in a duration- and dose-dependent manner. For instance, the use of a two-inhibitor (2i: PD0325901 and CHIR99021) cocktail, which chemically inhibits MEK/ ERK and GSK3 signaling respectively, has been instrumental in the reprogramming, maintenance, and derivation of naïve pluripotent stem cells (Nichols et al., 2009; Nichols and Smith, 2009; Silva et al., 2008; Ying et al., 2008). This approach has led to the acquisition of a stable and relatively homogeneous naïve pluripotent stem cell identity in culture, reflecting the naïve pluripotent cell compartment of the preimplantation embryo. Intriguingly, the small molecules CHIR99021 (a GSK3 inhibitor) and E616452 (Repsox, an ALK5 inhibitor) have frequently been used for the direct induction of many different cell types, e.g. chemical induced pluripotent stem cells (Hou et al., 2013; Zhao et al., 2018), cardiomyocytes (Cao et al., 2016; Fu et al., 2015) and neural progenitor cells (Cheng et al., 2014). Furthermore, retinoic acid signaling via a retinoic acid receptor (RAR) agonist has been shown to improve not only the reprograming of somatic cells into iPSCs (Wang et al., 2011); but also to induce marker expression and characteristics associated with totipotency (Hu et al., 2023; Iturbide et al., 2021; Wang et al., 2011; Xu et al., 2022).

Here, we employed insights from reprogramming to find small-molecule compounds capable of reprogramming pluripotent stem cells into founder-like precursor cells capable to directly generate a whole embryo model. This led to the generation of a chemical-only, transgene-free, cell mixing-free, direct, rapid, and efficient embryo model that recapitulates development into late streak stage with high molecular and morphological fidelity. These findings have significant implications not only for the development of embryo models but also for a variety of other biological research fields, owing to their underlying biology and ease of application.

## Results

### Chemical-only screen reprograms ESCs into putative precursors of blastocyst fates

Mouse extra-embryonic and embryonic lineages arise during the first and second embryo lineage specification, driven by the expression of key transcription factors such as *Cdx2*, *Gata6,* and *Oct4* (Mihajlovic and Bruce, 2017). These antagonistic factors are also co-expressed at the very early stages of embryo development at a time when lineage commitment hasn’t yet occurred, making them excellent reporters to attempt the generation of embryo founder-like/ precursor cells capable of giving rise to all early embryo fates. To generate such cells, we employed a candidate chemical-only screen and analyzed their ability to induce *Cdx2* and *Gata6* expression in ESCs while maintaining *Oct4* expression. We started with the medium formulation employed to generate morula-like cells (MLCs), which contains GSK3β inhibitor CHIR99021 (C) and mLIF (L) (Li et al., 2024; Li et al., 2023). However, unlike for the generation of MLCs, no reprogramming transgene was used in the candidate chemical screen presented here. As potential candidates, we considered small molecule compounds known to support the chemical-only generation of iPSCs (Hou et al., 2013; Zhao et al., 2018), which have been demonstrated to also produce primitive endoderm (PrE) fate cells. We hypothesized in our study that, instead, these molecules might underlie the generation of “plastic” progenitors capable of giving rise to all blastocyst fates.Interestingly, treating ESCs for 60 hours (h) with two of such small molecules, 6165452 (6) and AM580 (A), combined with CL (CL6A), resulted in robust activation of GATA6, minor activation of CDX2, while maintaining OCT4 expression (**Figure S1A**). However, since we also aimed to induce robust CDX2 expression to increase the likelihood of generating cells capable of developing into trophectoderm, we refined our formulation strategy. To enhance CDX2 expression, we introduced a preparing step for the first 24 hours before applying CL6A. We adjusted two variables, mLIF removal and the response to different concentrations of GSK3β inhibitor (C, 3 μM or C^+^, 6 μM or C^++^, 9 μM) (**Figure 1A and S1A**). Increasing concentrations of CHIR99021 has been previously shown to modulate fate decisions, but not yet been demonstrated for trophectoderm fate (Kempf et al., 2016; Yang et al., 2023). Removing mLif during the first 24 hours (cocktail K2) slightly increased *Cdx2* expression (**Figure S1A**). By increasing the concentration of CHIR99021 (K4 cocktail) during the first 24 hours and then treating with CL6A for 36 hours, we significantly increased the yield of cells with *Cdx2* expression from 1.28% to 29.2% of cells (**Figure 1B, 1C and S1A**). Additionally, Gata6 expression was detected in 27% of cells, while 77% cells maintained high OCT4 expression (**Figure 1C and S1A**). Immunostaining for CDX2 and GATA6 confirmed the trends of our K1-K4 chemical treatments (**Figure S1A and S1B**). qPCR analysis further validated that removing of mLIF for the first 24 hours and then increasing the concentration of CHIR99021 for the following 36 hours dramatically enhanced the induction of early TE associated transcription factors, e.g. *Cdx2*, *Gata3*, and *Id2*; and of ICM/ uncommitted-PrE associated markers (*Gata6*, *Pdgfr*α and *Sox17*) (**Figure S1C**).

**Figure 1.**
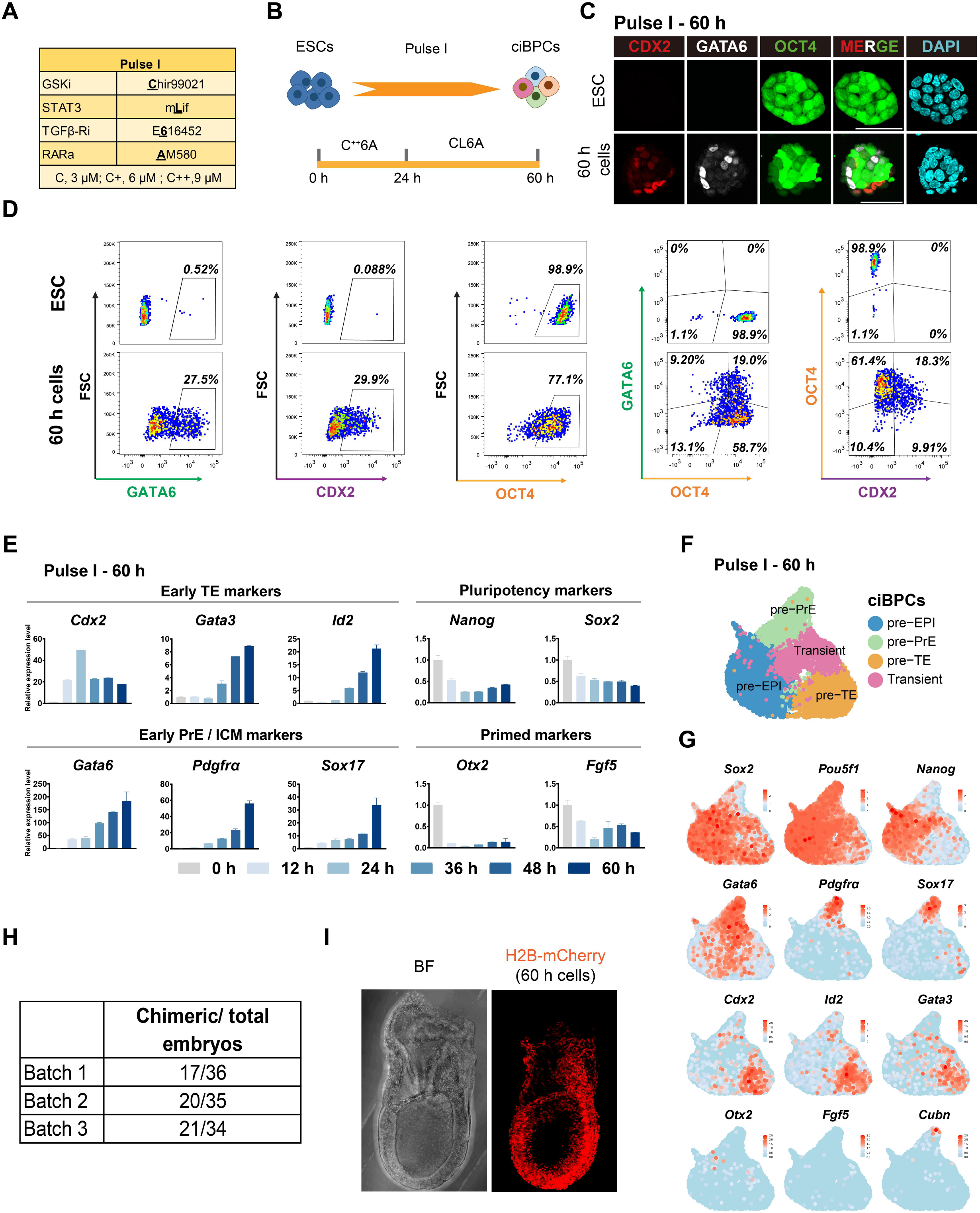
Screen identifies molecules that induce the generation of blastocyst-like precursor cells. See also Figure S1. (A) Small molecules referred to in this article and associated abbreviations are noted. C (Chir99021): glycogen synthase kinase 3 inhibitor (GSK3i); concentrations 3 μM (C), 6 μM (C^+^), 9 μM (C^++^). L (mLIF) represents mouse leukemia inhibitory factor, which activates JAK and STAT3 signaling. 6 (E616452) represents the selective transforming growth factor β receptor I inhibitor (TGFβ-Ri). A (AM580) represents the retinoic acid nuclear receptor (RAR) agonists. (B) Schematic of small molecules induction (K4 treatment) strategy. (C) Immunofluorescence of 60 h (pulse I) induction cells for TE marker CDX2 (red) and PrE marker GATA6 (white). ICM marker OCT4 (green) corresponds to the GFP signal driven by endogenous Oct4 activity; nuclear marker DAPI (cyan). Scale bar, 50 μm. (D) Flow cytometry for GATA6, CDX2, and OCT4 in 60 h (pulse I) induction cells. (E) Relative expression of marker genes in 60 h (pulse I) induction cells at different time points. Fold changes in mRNA expression of target genes were determined by the ΔΔCt method using *Actin* as endogenous control. (F) Single-cell sequencing UMAP of pulse I induction cells. Green represents “pre-PrE” cluster; orange represents “pre-TE” cluster; blue represents “pre-EPI” cluster; pink represents the “Transient” cluster. (G) Representative markers of different cell lineages. *Sox2*, *Pou5f1 and Nanog* mark EPI.; *Gata6*, *Pdgfr*α and *Sox17* mark PrE; *Cdx2*, *Id2* and *Gata3* mark TE; *Otx2* and *Fgf5* mark mature EPI; *Cubn* marks mature PrE. (H) Chimera formation ratio of 60 h induction cells. (I) Representative image of chimera from 60 h induction cells (H2B-mcherry, red).

Comparatively, the K4 chemical treatment is more effective at inducing both *Gata6* and *Cdx2* expression than K1-K3 treatments (**Figure S1A and 1C**). Therefore, we focused on the K4 cocktail, which involves treating ESCs with C^++^6A for 24 hours, followed by CL6A for 36 hours (**Figure 1B).** Interestingly, among the K4-treated cells at 60 h, we observed that 19% of cells co-express GATA6 and OCT4; and 18% of cells co-express CDX2 and OCT4 (**Figure 1D**). These results suggest that K4-treated cells acquired molecular signatures characteristic of earlier development stages, rather than specifying the fates of embryonic day (E) 4.5 blastocyst. qPCR time course analysis of K4 induction showed that early TE genes (*Cdx2*, *Gata3*, and *Id2*) and early ICM/ uncommitted pre-associated genes (*Gata6*, *Pdgfr*α, and *Sox17*) were activated, whereas expression of naïve pluripotency-associated genes (*Nanog* and *Sox2*) slightly decreased (**Figure 1E**). Importantly, early differentiating genes *Fgf5* and *Otx2* did not show increased expression, further suggesting that the observed biological changes are not related to ESC developmental progression but rather indicate possible reprogramming to earlier developmental stages (**Figure 1E**).

To further investigate the cellular fates of chemically induced cells, we performed scRNA-seq on cells at 60 h time point (**C^++^6A + CL6A/ pulse I**) using 10x Genomic Chromium. When projected onto a uniform manifold approximation and projection (UMAP), four clusters could be defined based on molecular signatures in 60 h cells: Epiblast precursor cells (**pre-EPI**), PrE precursor cells (**pre-PrE**), TE precursor cells (**pre-TE**), and transient-state cells (**Transient**), a cluster comprised of potentially less biased subpopulations (**Figure 1F**). At the single-cell level, we observed the co-expression of *Pou5f1* and *Gata6* and of *Pou5f1* with *Cdx2* (**Figure 1G**). Markers indicating commitment such as *Otx2*, *Fgf5* and *Cubn* were barely expressed in these cells.

We also investigated the reproducibility of our findings by using two additional independent ESC lines derived from different mouse strains, B6 and ICR ESC. Immunostaining for key factors (CDX2/ GATA6/ SOX2) consistently validated our results across all three lines used (**Figure S1D and S1E**).

As these cells were exposed to novel culture conditions, we tested their ability to enter embryo development in a chimera assay. We injected 4-6 cells labeled with H2B-mCherry into 8-cell embryos. Chimerism was examined at E7.5, and different batches showed that more than half of the embryos displayed a clear contribution of 60 h cells (**Figure 1G**). Additionally, the mCherry signal revealed high chimerism (**Figure 1H**) demonstrating that 60 h cells possess robust ability to enter development.

These findings show that we identified small-molecule compounds that successfully induce the expression of *Cdx2* and *Gata6* while maintaining *Pou5f1* expression, resulting in the generation of putative precursors of blastocyst-like fates.

### Committing precursor cells into late blastocyst Fates

Although the expression of early lineage markers CDX2 and GATA6 was induced, markers associated with lineage commitment at the late blastocyst stage, such as GATA4 protein (Suzuki et al., 2022a) and *Cubn* mRNA (Kurimoto et al., 2006), were barely detected in 60 h cells. To promote cell fate specification into E4.5-like Epiblast (EPI), PrE, and TE identities, we introduced a second pulse (pulse II) of small molecule treatment. Several defined culture conditions for inducing TSCs and XEN cells contain the same molecules, e.g. FGF4, and BMP4 (Artus et al., 2012; Seong et al., 2022). We also investigated if LATS inhibitor TRULI (Gao et al., 2024; Nishioka et al., 2009) could enable the 60 h cells to undergo E4.5 TE fate specification. To promote 60 h cells to adopt E4.5 epiblast identity, we added ACTIVIN A and XAV939, also used in media to induce trophectoderm fate (Gao et al., 2024; Nishioka et al., 2009). Through medium optimization, we defined the following set of molecules: **<u>T</u>**RULI (2 μM), ACTI**<u>V</u>**IN A (20 ng/mL), **<u>X</u>**AV939 (3 μM), **<u>B</u>**MP4 (10 ng/mL), and **<u>F</u>**GF4 (25 ng/mL), **TVXBF**, as inducers of 60 h precursors into committed blastocyst fates (**Figure 2A**). This medium is hereafter defined as **Priming medium (PM)**. To test this medium formulation, we dissociated the 60 h cells (pulse I) and exposed them to PM for 36 hours in Aggrewell at a density of 25∼30 cells per microwell (**Figure 2A**). Maturation into the PrE fate was confirmed by the increased expression of GATA4 in 96 h cells compared to the 60 h cells (**Figure 2A**). Morphologically, the GATA4^+^ cells were found at the boundary of 96 h cell clusters, which is feature of primitive endoderm-like cells (**Figure 2B**). The skeleton proteins KRT18 and cell surface marker PDGFRα further support that 96 h cells adopted E4.5 blastocyst-like identities (**Figure 2C**).

**Figure 2.**
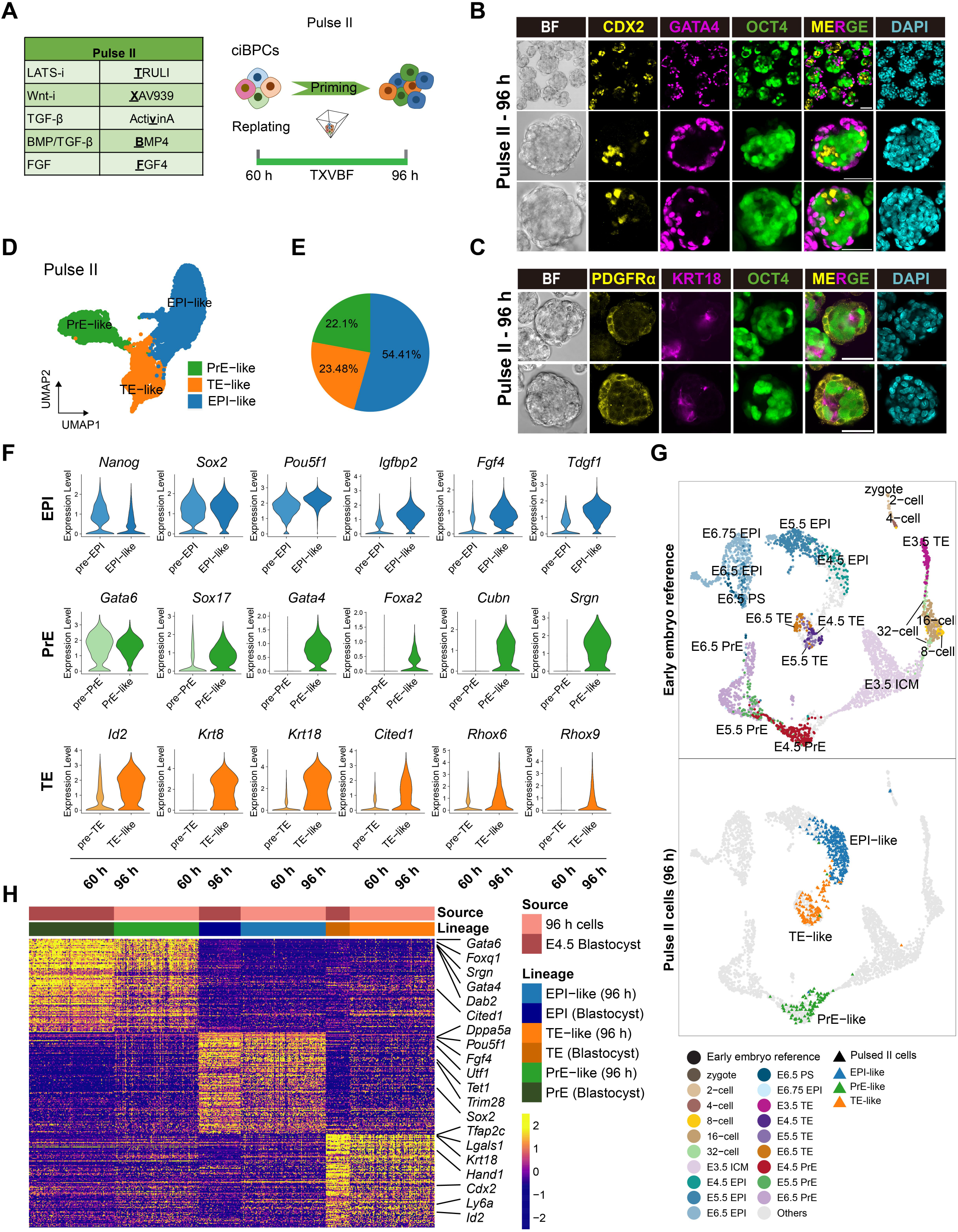
Trajectory analysis of chemically induced cells reveals pathways to blastocyst-like lineages. See also Figure S2. (A) Schematic of Pulse II small molecules induction strategy. T (TRULI) represents the LATS inhibitor. X (XAV939) represents the Wnt signaling inhibitor. V (ActivinA) is one of the TGFβ superfamily members. B (BMP4), TGFβ family. F (FGF4), a member of the fibroblast growth factor (FGF) family. (B-C) Immunofluorescence of 96 h (pulse II) induction cells. TE marker CDX2 (yellow), KRT18 (purple); PrE marker GATA4 (red), PDGRFα (yellow). ICM marker OCT4 (green) corresponds to the GFP signal driven by endogenous Oct4 activity; nuclear marker DAPI (cyan). Scale bar, 50 μm. (D) Single-cell sequencing UMAP of pulse II induction cells. Green represents PrE-like cluster, orange represents TE-like cluster, blue represents EPI-like cluster. (E) Statistical proportions of the three clusters for pulse II induction cells. (F) Representative markers of different cell lineages. *Nanog*, *Sox2*, *Pou5f1*, *Igfbp2*, *Fgf4* and *Tdgf1* mark EPI.; *Gata6*, *Gata4*, *Sox17*, *Foxa2*, *Cubn* and *Srgn* mark PrE; *Cited1*, *Rhox6*, *Rhox9*, *Id2*, *Krt8* and *Krt18* mark TE. (G) UMAP of 96 h induction cells integrating with published datasets from early mouse embryo stages from zygote till E6.5 (Cheng et al., 2019; Deng et al., 2014; Mohammed et al., 2017; Nowotschin et al., 2019; Posfai et al., 2017). (H) Heatmap showing the expression levels of conserved genes among 96 h cells and E4.5 blastocysts.

Strikingly, at 96 h, cells were organized into three distinct clusters with a ratio of 22.1% (PrE-like), 23.48% (TE-like), and 54.41% (EPI-like) each showing robust expression of respective commitment markers for E4.5 PrE (*Gata4*, *Sox17*, *Gata4*, *Foxa2*, *Cubn*, and *Srgn*), E4.5 TE (*Cited1*, *Id2*, *Krt8*, *Krt18*, *Rhox6,* and *Rhox9*) and of E4.5 EPI (*Fgf4*, *Tdgf1*, *Igfbp2,* and *Tdgf1*) respectively (**Figure 2D-2F**).

Next, we performed dimension reduction analysis using UMAP against scRNA-seq data (96 h) from zygote till E6.5 (Cheng et al., 2019; Deng et al., 2014; Mohammed et al., 2017; Nowotschin et al., 2019; Posfai et al., 2017) (**Figure 2G**). The PrE-like, TE-like, and EPI-like clusters were perfectly mapped with E4.5 blastocyst PrE, TE, and EPI cells respectively (**Figure 2G**). Furthermore, we explored the expression levels of top lineage markers of E4.5 blastocyst fates in both populations to evaluate the similarities between 96 h cells and E4.5 blastocyst fates. Importantly, these display a comparable expression pattern, meaning that EPI-, TE- and PrE-like cells are transcriptionally similar to embryo E4.5 blastocyst fates (**Figure 2H and Table S1**).

Together, these data show that the second pulse (pulse II) of small molecule compounds promoted the specification of embryo founder-like/ precursor cells into committed blastocyst fates.

### Fate and trajectory analysis of chemically induced ESCs

To get a better understanding of the trajectory leading to the generation of blastocyst-like precursor cells we performed refined clustering analysis at 24 h, 60 h, and 96 h time points (**Figure 3A**). At 24 hours, we could already observe up-regulation of *Cdx2* and *Gata6* expression, indicating that reprogramming starts immediately after chemical treatment (**Figure 3A and 3B**). By 60 hours, cells displayed an up-regulation in either E3.5 TE or E3.5 ICM scores, suggesting that reprogramming towards a state resembling an earlier developmental stage relative to E4.5 EPI-like starting ESCs is occurring (**Figure 3A and 3C)**. Reprogramming towards E3.5 ICM and to E3.5 TE-like state in 60 h (pulse I) cells was confirmed by the activation of *Calcoco2 (Spangler et al., 2018)*, *Khdc3 (Boroviak et al., 2018)*, and *Bhmt* (Lee et al., 2012; McClatchie et al., 2017), as well as by *Crb3 (Jiang et al., 2018), Cldn7* (Moriwaki et al., 2007) and *Cdx2* (Suzuki et al., 2022b) respectively (**Figure 3C**). In 96 h (pulse II) cells, clear lineage segregation into E4.5 blastocyst stage fates is observed (**Figure 3A and 3D**).

**Figure 3.**
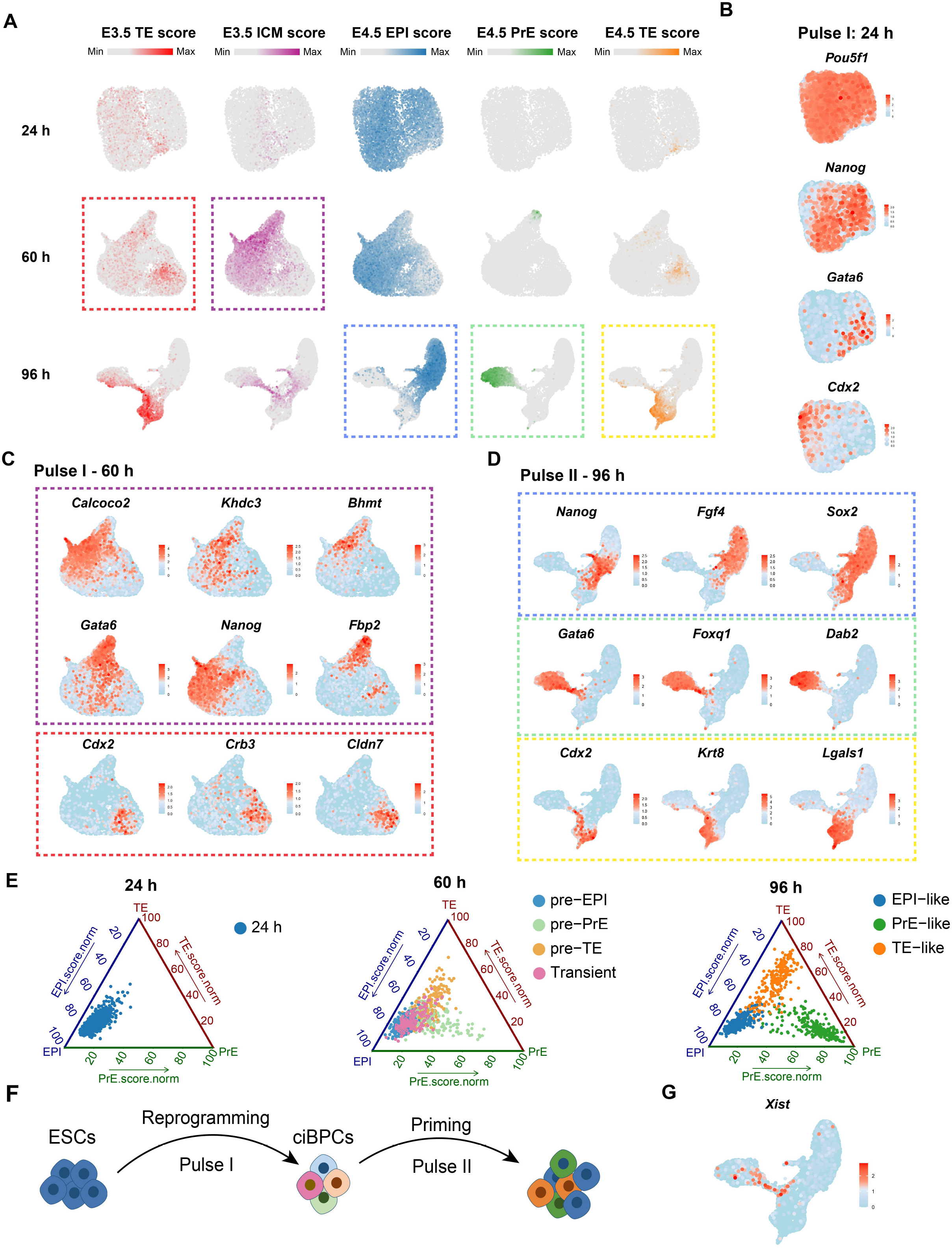
Chemically induced ESCs undergo reprogramming before committing to specific blastocyst fates. (A) Normalized lineage scores for 24 h, 60 h, and 96 h induction cells calculated using the top 50 differentially expressed genes (|Log2 Fold Change| >1 and FDR < 0.05) of lineages of E3.5 (E3.5 TE scores in red and ICM scores in purple) and E4.5 embryos (EPI, PrE and TE score are indicated in blue, green and orange respectively). The scores were normalized across all cells from all induction time points. (B) Expression levels of key transcription factors *Pou5f1*, *Nanog*, *Cdx2*, *Gata6* of blastocyst lineages at 24 h cells of pulse I. (C) Expression levels of selected representative differentially expressed genes of E3.5 ICM (framed in purple) and TE cells (framed in red) used in the computation of lineage scores at 60 h cells of pulse I. (D) Expression levels of selected representative differentially expressed genes of E4.5 EPI (framed in blue), PrE (framed in green), and TE (framed in orange) cells used in the computation of lineage scores at 96 h cells of pulse II. (E) Lineage identity was further confirmed by lineage score analysis. The ggtern plots show the normalized lineage scores for each individual cell, calculated by using the top 50 differentially expressed genes (|Log2 Fold Change| >1 and FDR < 0.05) of lineages of E4.5 embryos. The scores were normalized using the score of E4.5 embryos cells as maximum value. Higher values indicate closer similarity to E4.5 natural embryos. The cells are colored according to their lineage identity. (F) Schematic of small molecules induction strategy. (G) Expression levels of *Xist* in 96 h XY cells.

To decipher when development towards committed E4.5 blastocyst lineage takes place, we defined the three E4.5 blastocyst counterparts as having a 100% scoring for either EPI, PrE and TE similarity, and measured the trajectory of 24 h, 60 h and 96 h cells (**Figure 3E and Table S2**). Compared with the 24 h E4.5 EPI-like starting cells, the 60 h extra-embryonic lineages precursor cells showed slightly decreased EPI similarity while gradually acquiring the characteristics of either PrE or TE lineages as was indicated by the minor up-regulation of corresponding similarity scores. Combined with the up-regulation of E3.5 embryo similarity at 60 hours, these again suggest a plastic/ transient state by 60 h. A sharp increase in the three scores took place in 96 h compared with 60 h cells. Combined with the previously mentioned up-regulation of mature lineage markers (**Figure 2F**), these indicated the commitment towards the E4.5 blastocyst lineages (**Figure 3E and Table S2**).

Of biological interest, we observed transient upregulation of *Xist* expression in male XY cells at the intersections of EPI, PrE and TE fates (**Figure 3G**). This not only indicates exit from a naïve pluripotent cell identity (Sousa et al., 2018) but also suggests that TE and PrE fates in our system can regulate *Xist* expression, even though *Xist* regulation in extraembryonic lineages is pre-determined by imprinting during natural embryo development.

Taken together, these findings reveal that cells undergo reprogramming up to the 60 h time point. Following the second pulse of chemical treatment, they then develop into one of the late blastocyst fates (**Figure 3F**).

### 96 h TE-like cells show mural and polar specification

In the late blastocyst, the TE lineage undergoes further development, with mural TE characterized by increased expression of laminins, integrins, and ephrins (Klaffky et al., 2001; Sutherland, 2003). To investigate whether 96 h TE cells are comparable with either polar or mural TE, we focused our analysis on the TE-like cells from the scRNA-seq data. Interestingly, we found that these cells expressed polar TE markers (*Cdx2*, *Ddah1*, *Ly6a*, *Gsto1* and hippo pathway target *Ccnd1*) and/ or mural TE markers (such as intermediate filaments *Krt18*, *Krt8*, integrin *Itga6*, galectin *Lgals1*, and ephrin *Efnb1*) (Seong et al., 2022) (**Figure S2A and S2B**). Based on the expression patterns, we classified the TE sub-clusters into mural TE-like (cluster 1 and cluster 4) and polar TE-like (cluster 0, cluster 2 and cluster 3). In order to verify the cell identities, we computed the differentially expressed genes of each cell cluster and used them for the Gene Ontology (GO) enrichment and Kyoto Encyclopedia of Genes and Genomes (KEGG) pathway enrichment for the sub-clusters of the TE-like cluster (**Figure S2C, S2D and Table S3**). As expected, all TE-like sub-clusters exhibited enrichment of trophectodermal cell fate-related terms such as “Trophectodermal cell differentiation” and “Placenta development” to varying degrees. Additionally, cluster 1 and cluster 4 showed enrichment in genes associated with “Actin filament organization”, strongly suggesting the functional characteristics of mural TE (**Figure S2C**). KEGG pathway enrichment analysis showed that putative mural cell clusters (cluster 1 and cluster 4) are enriched for “Focal adhesion” and “Tight junction” pathways (**Figure S2D**), which are characteristic of embryo mural TE at the initial stage of implantation. In the late mouse blastocyst, the CDX2 marks the polar TE, while KRT18 increases in the mural TE of the E4.5 blastocyst. Also, the immunostaining for KRT18 and CDX2 confirmed differing degrees of expression in the polar TE-like cells and mural TE-like cells in 96 h cells (**Figure S2E**). Taken all these evidence together, we propose that 96 h TE-like cells derived from chemically treated ESCs faithfully recapitulate the development of both polar and mural TE in the late blastocyst, as indicated by specific marker expression and pathway enrichment profiles.

### ci-MOs recapitulate embryo development

The 96 h cells, which encompass all the cell types that characterize the blastocyst, were therefore hypothesized to have a high potential for developing into an embryo model, referred to as chemical-induced embryo model (ci-MO). The 96 h cells, now defined as day (d) 0 ci-MOs, were used to test their competence to generate embryo-like structures using the FC-IVC1-IVC2 culture system, known to support post-implantation embryo model development (Amadei et al., 2022; Lau et al., 2022). In FC medium, ci-MOs compacted and arranged into three distinct cell compartments within 1.5 days, marked by CDX2^+^/ AP2γ^+^ (TE-origin), OCT4^+^ (EPI-origin) or GATA6^+^/ GATA4^+^ (PrE-origin) (**Figure 4A, S3A, and S3B**). The PrE-derived compartment surrounded the EPI and TE compartments, similar to the post-implantation embryo. These well-organized structures constitute 85% of all structures recovered from any given well. After 1.5 days, the FC medium was replaced with the IVC1 medium. By day 2.5, ci-MOs displayed rosette-to-lumen formation in both EPI- and Extraembryonic ectoderm (ExE)-like compartments (**Figure 4A, S3C, and S3D**), resembling E5.0∼5.5 embryos (Langkabel et al., 2021). The efficiency of ci-MOs undergoing rosette-to-lumen formation from day 1.5 to day 2.5 was on average 70%. Due to the increased size, day 2.5 ci-MOs were transferred to 6-well non-adherent tissue culture plates in a shaking incubator at 80 rpm for 1 day. By day 3.5, ci-MOs elongated and developed egg-cylinder structures that closely resemble E6.5 natural embryos. These consist of EPI (SOX2, OCT4) and ExE (SOX2, AP2γ) compartments, both enveloped by a visceral endoderm-like structure (**Figure 4A, S3E**). The EPI displayed an expanded pro-amniotic cavity and successfully broke symmetry, indicating the onset of the primitive streak on one side of the EPI, adjacent to the ExE. The establishment of the anterior-posterior axis is critical for proper development (Belo et al., 1997; Thomas and Beddington, 1996). Following the onset of gastrulation in the posterior EPI, the primitive streak (T/ Bry) extends to the distal end (**Figure 4A**). The efficiency of ci-MO development till the onset of gastrulation, from day 2.5 to day 3.5, was on average over 70%. We also investigated the reproducibility of our findings using two additional independent ESC lines derived from different mouse strains, B6 and ICR (**Figure S4A and S4B**). Again, our results were consistently validated across the three lines used. These results show that ci-MOs successfully form distinct embryo-like compartments and develop into advanced stages resembling natural embryos.

**Figure 4.**
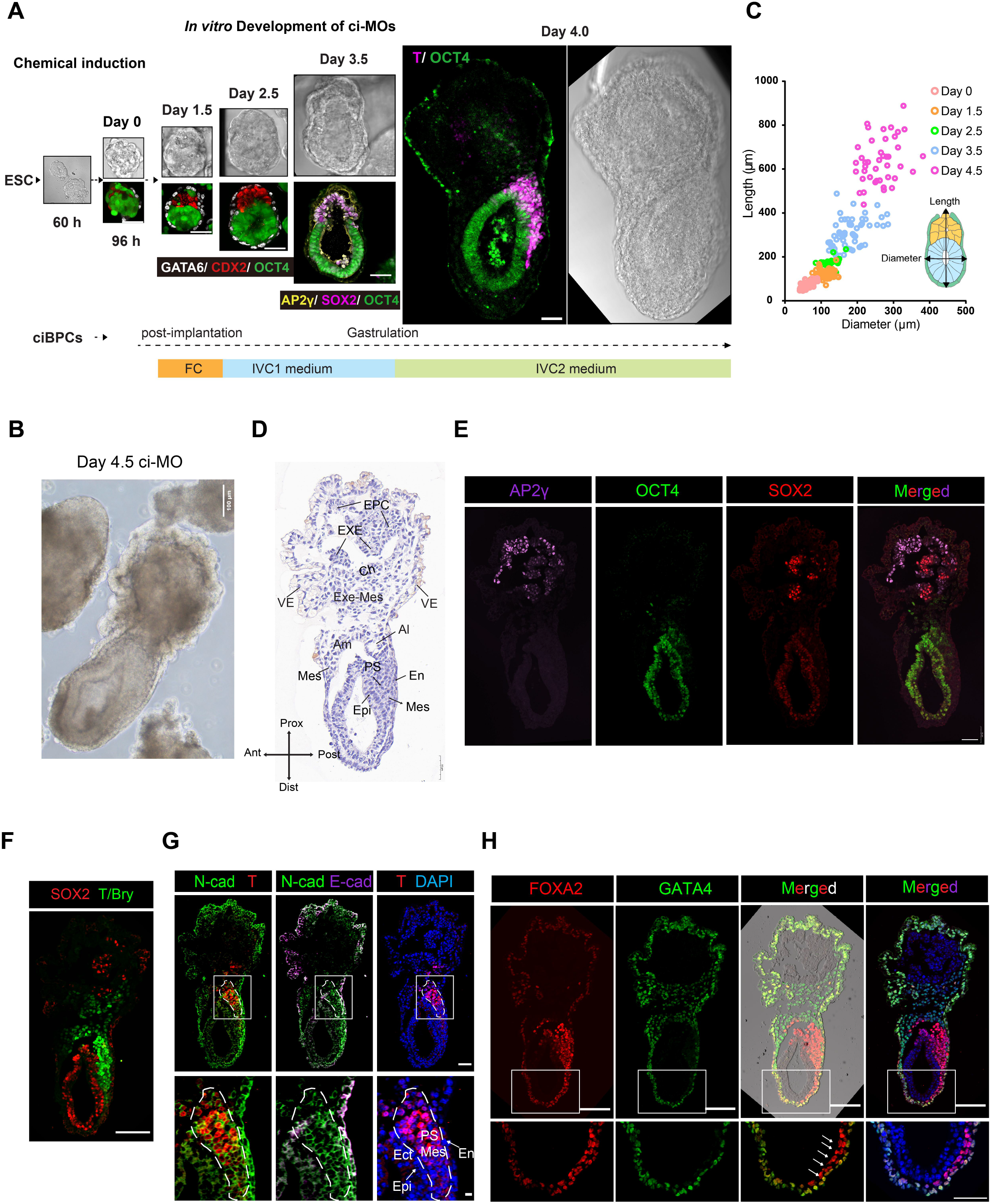
Formation and development of ci-MOs from 96 hour chemically induced ESCs. See also Figure S3 and S4. (A) Progression of ci-MOs from ESC to 96 h ciBPCs till day 4.0 ci-MOs with representative staining *in vitro*. Scale bar, 100 μm. (B) Morphology of day 4.5 ci-MOs. Scale bar, 100 μm. (C) Length and diameter change with ci MOs progression. (D) Paraffin of day 4.5 ci-MOs. EPC, ectoplacental cone; EXE, extraembryonic ectoderm; Ch, chorion; Exe-Mes, extraembryonic mesoderm; VE, visceral endoderm; Am, amnion; Al, allantois; Mes, mesoderm; PS, primitive streak; En, endoderm; Epi, epiblast; Prox, proximal; Dist, distal; Ant, anterior; Post, posterior. Scale bar, 50 μm. (E-H) Immunofluorescence of day 4.5 ci-MOs paraffin. F, day 4.5 ci-MOs analyzed for AP2γ (purple), OCT4 (green), SOX2 (red), Scale bar, 50 μm. G, day 4.5 ci-MOs analyzed for SOX2 (red) and T (green), Scale bar, 50 μm. H, day 4.5 ci-MOs analyzed for N-cad (green), T (red), E-cad (purple), and DAPI (blue); Mes, mesoderm; PS, primitive streak; En, endoderm; Epi, epiblast; Ect, ectoderm; Scale bar, 50 μm. I, day 4.5 ci-MOs analyzed for FOXA2 (red), GATA4 (green), DAPI (blue), Scale bar, 50 μm.

Next, we evaluated the generation of ci-MOs both quantitatively and qualitatively, focusing on size and molecular signatures. Over 4.5 days, ci-MO development was marked by changes in size, as indicated by measured diameter and length (**Figure 4A and 4B**). Additionally, the embryo morphology changes from day 1.5-2.5 to day 4.0 ci-MOs closely followed the development of embryos from E5.5 to gastrulation stages (**Figure 4A**). By day 4.5, ci-MOs exhibited high morphological and molecular similarity to E7.5 embryos (**Figure 4C-4H**). At the transition from day 3.5 to day 4.5, the proportion of structures exhibiting gastrulating embryo morphologies exceeded 37%. Histological section staining of day 4.5 ci-MO confirmed the presence of various embryonic and extraembryonic tissues and structures similar to an E7.5 mouse embryo (**Figure 4D**). Notably, we observed that the gastrulating ci-MOs recapitulated the segregation of germ-layer tissues (epiblast, endoderm, and mesoderm), epithelial to mesenchymal transition (EMT), formation of the visceral endoderm and definitive endoderm, ectoplacental cone (EPC), allantois (Al), and amnion (Am) (**Figure 4D**). Accordingly, day 4.5 ci-MOs displayed the differentiation of TE into ExE (SOX2^+^ weaker AP2γ^+^ OCT4^-^ counterpart) and EPC (Bright AP2γ^+^) (**Figure 4E**). Primitive streak emerged from EPI and extended into the distal portion (**Figure 4F).** EMT was evidenced by the downregulation of E-Cadherin and the upregulation of N-Cadherin (**Figure 4G**), a key event in gastrulation. We also observed the presence of FOXA2 positive cells overlapping part of the PS (**Figure 4F and 4G**), which defines axial mesendoderm specification (Arnold and Robertson, 2009). Meanwhile, FOXA2-positive and GATA4-negative cells marked definitive endoderm, seen here gradually intercalating with the VE (FOXA2^+^, GATA4^+^) cells enveloping the egg cylinder (**Figure 4H**).

These findings demonstrate that ci-MOs not only mimic the morphological and molecular characteristics of natural embryo development from E4.5 to E7.5 but also successfully recapitulate critical gastrulation events and tissue differentiation.

### Day 4.5 ci-MOs display high-fidelity E7.5 molecular signature

Day 4.5 ci-MOs replicate numerous gastrulation events, so we sought to compare them to natural embryos to determine the corresponding embryo developmental stage and overall cell type composition. We performed scRNA sequencing on pooled day 4.5 ci-MOs (n=4, with gastrulating morphology) using 10x Genomics Chromium. After stringent quality control to filter out low-quality cells, we obtained 15441 single-cell transcriptomes with a unique molecular identifier (UMI) of 4058 genes detected per cell. The median mitochondrial read portion was 1.15%, suggesting good cell quality. The embryo reference scRNA sequencing dataset that was shown to match closer to day 4.5 ci-MOs was the E7.5 natural embryo (**Figure 5**), capturing mouse gastrulation (Bardot and Hadjantonakis, 2020; Pijuan-Sala et al., 2019).

**Figure 5.**
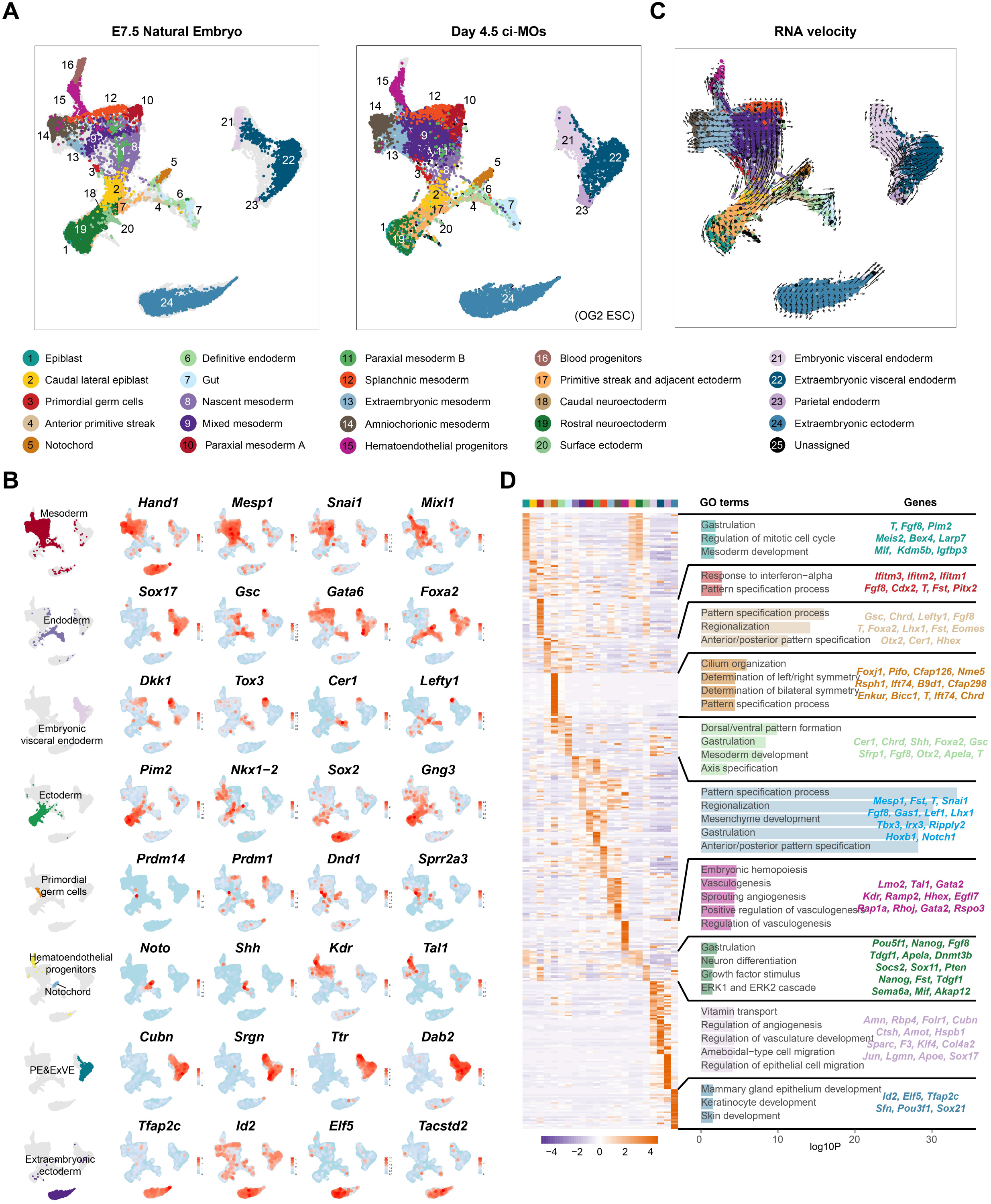
Single-cell analysis of day 4.5 ci-MOs. See also Figure S5. (A) A Uniform Manifold Approximation and Projection (UMAP) analysis of scRNA-seq data from E7.5 natural embryos (public) and day 4.5 ci-MOs (this study). (B) UMAP plots showing expression of indicated lineage markers in day 6 ci-MOs. *Hand1*, *Mesp1*, *Snai1*, and *Mixl1* mark Mesoderm; *Sox17*, *Gsc*, *Gata6*, and *Foxa2* mark Endoderm; *Dkk1, Tox3, Cer1, and Lefty1* mark anterior visceral endoderm; *Noto* and *Shh* mark Notochord; *Kdr*, and *Tal1* mark hematopoietic endothelial progenitor cells; *Pim2*, *Nkx1-2*, *Sox2*, and *Gng3* mark Ectoderm; *Prdm14*, *Prdm1*, *Dnd1*, and *Sprr2a3* mark Primordial germ cells; *Cubn*, *Srgn*, *Ttr*, and *Dab2* mark Parietal endoderm and Extraembryonic visceral endoderm (PE & ExVE); *Tfap2c*, *Id2*, *Elf5*, and *Tacstd2* mark Extraembryonic ectoderm. (C) The RNA velocity field is projected onto the UMAP plot shown in the right panel A. (D) Heatmap of expression patterns of the top 30 differential expressed genes (log2FC > 1) per cell type (left), enriched GO terms (middle), and representative genes associated with each GO term (right).

Clustering analysis based on differentially expressed genes revealed 24 different cell identities (**Figure 5A**). The annotation of these identities was accomplished based on the major cell lineage previously defined by single-cell transcriptomics of mouse embryos (Qiu et al., 2022). Strikingly, the profile of cell types found in day 4.5 ci-MOs was highly similar to E7.5 natural embryo, demonstrating that lineage commitment and developmental complexity are faithfully recapitulated in ci-MOs at the single-cell molecular level (**Figure 5A**). All three germ layers were represented as well as all extraembryonic tissues, in a similar manner in both *in vivo* natural embryos and ci-MOs (**Figure 5A and 5B**).

Specifically, we identified gastrulation mesoderm-like lineage, characterized by the expression of genes such as *Hand1 (Barnes et al., 2010)*, *Mesp1 (Ajima et al., 2021)*, *Snai1*, and *Mixl1(Lindsley et al., 2006)* (**Figure 5A and 5B**). Consistent with the immunofluorescent staining result from day 4.5 ci-MOs, we detected the presence of primitive streak characterized by *T*/*Bry* expression (**Figure 5A and Table S4**). Within the mesoderm cells, we identified several mesoderm lineages, including cluster 4, anterior primitive streak (*Gsc (Lewis et al., 2007)*, *Lefty (Takaoka et al., 2011)*, *T*, *Tdgf1*, *Hhex*, *Cer1*, *Eomes*); cluster 9, mixed mesoderm (*Mesp1, Crabp1, Foxf1*); cluster 8, nascent mesoderm (*Mesp1, Fst, Slpi*); cluster 10, paraxial mesoderm A (*Dll3, Ppp1r1a, Map1b*); cluster 11, paraxial mesoderm B (*Aldh1a2, Foxc1, Dll1*); cluster 12, splanchnic mesoderm (*Mef2c, Rbm24, Igfbpl1*); cluster 13: extraembryonic mesoderm (*Spin2c, Foxf1, Pmp22*); and cluster 14, amniochorionic mesoderm (*Tdo2, Dlk1, Ahnak*) (**Figure 5A and Table S4**). Extraembryonic mesoderm plays a role in early hematopoiesis and erythropoiesis by providing mechanical support to the developing embryo (Saykali et al., 2019). Adjacent to cluster 13, we identified a cell cluster signifying the hematoendothelial progenitors (cluster 15), marked by *Kdr* (Li et al., 2017), *Tal1* (Kallianpur et al., 1994) and *Pecam1* (Litwin et al., 1997), indicating the initiation of primitive hematopoiesis from extraembryonic mesoderm by RNA velocity (**Figure 5B and Table S4**). GO analysis confirmed the expression of several hematopoietic-related markers and enrichment in terms related to vasculogenesis and hematopoiesis (**Figure 5D, Table S4 and S5**).

We were able to detect ectoderm lineages in gastrulating ci-MOs, which are marked by *Nkx1-2 (Verrier et al., 2018)*, *Sox2 (Wood and Episkopou, 1999)*, *Gng3 (Kelly et al., 2008),* and *Pim2*, a newly defined ectoderm-specific marker in this study (**Figure 5A-4C**). Within the ectoderm cells (clusters), we identified post-implantation epiblast-like clusters: cluster 1, epiblast, which expressed core pluripotency transcription factor *Pou5f1* and *Nanog* and cluster 2, caudal lateral epiblast, which is characterized by the expression of genes such as *Fgf8*, *Pten*, *Kdm5b*, *Igfbp3* (**Figure 5A-5C**). Furthermore, we were able to validate the sub-clusters of ectoderm lineages: cluster 19, Rostral neuroectoderm, characterized by *Pim2, Gng3, Utf1*; cluster 20, Surface ectoderm, characterized by *Id1, Id3, Ifitm3*; and cluster 17, primitive streak and adjacent ectoderm, characterized by *Pim2*, *Fst* (**Figure 5A-5C**). We also detected definitive endoderm cells (cluster 6, *Foxa2*, *Gsc*, *Cer1*, *Dkk1*), and gut (cluster7, *Shh*), which is consistent with the immunofluorescent staining for FOXA2 results from day 4.5 ci-MOs (**Figure 4A and 4I**). Also, we identified the emergence of primordial germ cells (cluster 3), characterized by *Prdm14*, *Prdm1*, *Dnd1,* and *Sprr2a3*. Moreover, the distribution of cells expressing *Noto* and *Shh* marked the key event of Notochord (**Figure 5C-5D**).

After implantation, the primitive endoderm differentiates into visceral endoderm and parietal endoderm, which contribute to the formation of the yolk sac. Notably, we detected lineages with strong expression of *Cubn*, *Sox17*, *Ttr,* and *Dab2*, including cluster 21, embryonic visceral endoderm; cluster 22, extraembryonic visceral endoderm; cluster 23, parietal endoderm (**Figure 5B and 5D**). Gene enrichment analysis also revealed a distinct pattern involving angiogenesis and vasculature development. Remarkably, ci-MOs effectively mimic embryonic development.

When applied to an independent ESC line from a different mouse strain (B6), the day 4.5 ci-MOs successfully recapitulated the characteristics of E7.5 gastrulating embryos with high fidelity (**Figure S5 A-D, Table S6 and S7**).

The high similarity in lineage commitment and developmental complexity indicates that ci-MOs can faithfully recapitulate gastrulation events, making them robust models for studying embryonic development and cell fate specification.

### ci-MOs display advanced trophoblast lineage development

*In vivo*, shortly after implantation, the mural TE differentiates into polyploid trophoblast cells, which are also known as primary parietal trophoblast giant cells (primary P-TGCs) (Hannibal and Baker, 2016); while the undifferentiated polar TE forms the extraembryonic ectoderm (ExE). Trophoblast stem cells could be derived from both polar TE and ExE (Hayakawa et al., 2015; Tanaka et al., 1998); when the ectoplacental cone (EPC) arises from the continuing expansion of the ExE. The EPC cavity and the exocoelomic (chorionic) cavity form during gastrulation (E6.5-8.5) (Bardot and Hadjantonakis, 2020; Tam and Loebel, 2007).

To decipher TE lineage development in the gastrulating day 4.5 ci-MOs, we extracted the TE portion and compared it to a mouse placentation developmental roadmap from E7.5 to E14.5, a reference dataset revealing the murine placentation at single-cell resolution (Jiang et al., 2023) (**Figure 6**, Sub-clustering identified 7 clusters within day 4.5 ci-MOs TE: TE progenitors, TSC-1, TSC-2, EPC -1, EPC-2 (migratory EPC), decidualization-driving cells, and a cluster of exocoelomic (chorionic) cavity cells (marked by *Id2*, *Ttr*, *Sox17* and *Dab2*) (**Figure 6A**). RNA velocity results suggest that the TE progenitors cluster contains the progenitor cells of both TSC-1 and TSC-2, while EPC-2 cluster cells derive from EPC-1 cluster cell differentiation (**Figure 6B**). To our surprise, the EPC-2 cluster includes at least 6 subclusters of EPC migratory cells, including secondary mononucleated sinusoid trophoblast giant cells (S-TGC) precursors, secondary definitive placental TGC (P-TGC) precursors, syncytiotrophoblast cells (synT) precursors, labyrinth trophoblast progenitor cells (LaTP) cells and others (**Figure 6C**). These advanced cell types, which are developed at late-streak stage *in vivo*, not only suggest that the ci-MOs have both mural TE and polar TE origins(Rossant, 2018), but also indicate that the ci-MO system has advanced developmental potential in placenta derivatives (Pijuan-Sala et al., 2019).

**Figure 6.**
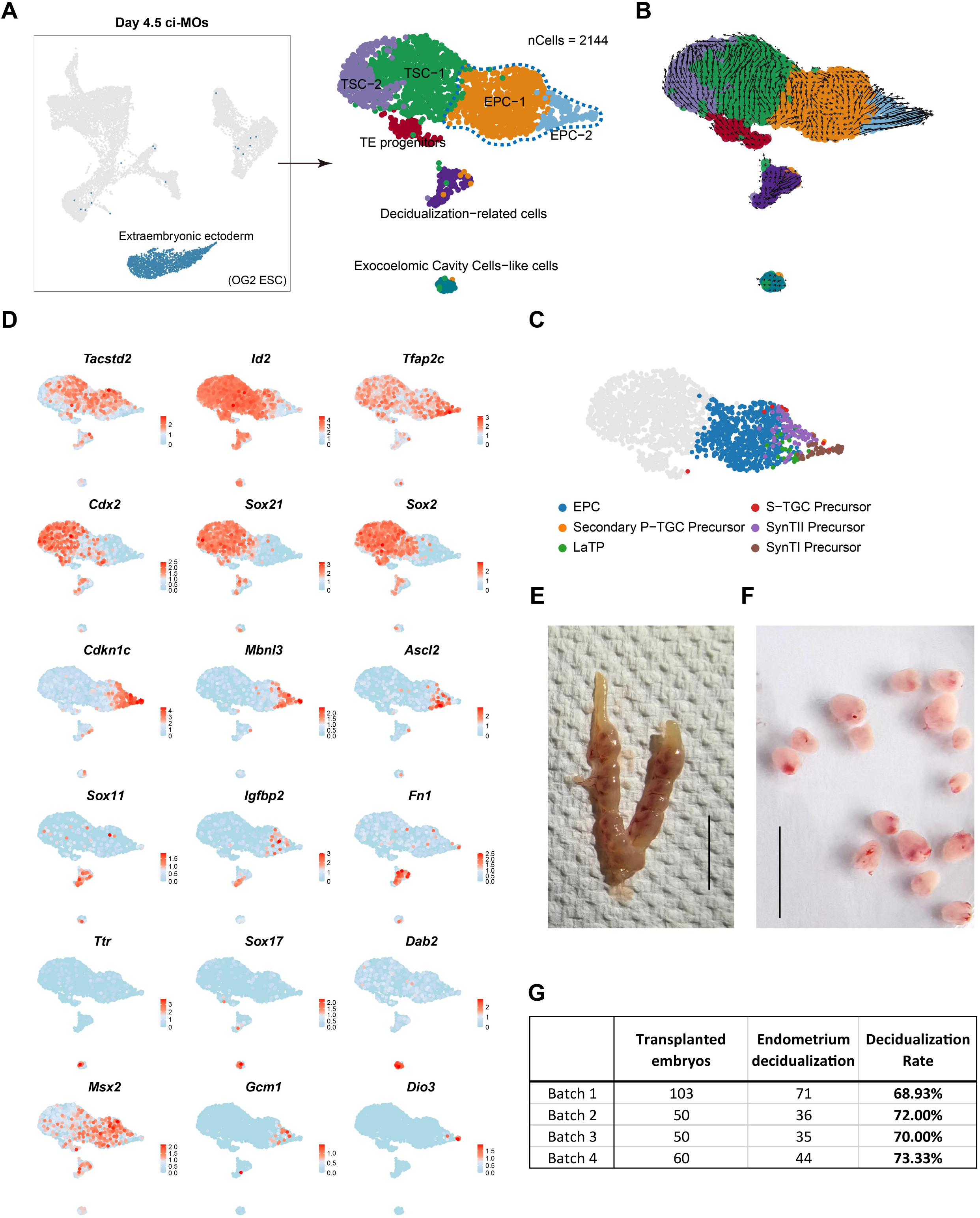
Characterization of extraembryonic ectoderm cell derivatives in day 4.5 ci-MOs. (A) Sub-clustering and annotation results of extraembryonic ectoderm cells using classical markers, and gene expression results in day 4.5 ci-MOs. (B) RNA velocity maps showing developmental trajectories of extraembryonic ectodermal cells. (C) Sub-clustering and annotation results of EPC clusters based on published datasets (Jiang et al., 2023). (EPC, ectoplacental cone; Secondary P-TGC precursor, Secondary primary parietal trophoblast giant cells; LaTP, labyrinth trophoblast progenitor cells; S-TGC Precursor, sinusoid trophoblast giant cells; SynTII/ I Precursor, syncytiotrophoblast precursor cells II/ I) (D) Heatmap of expression patterns of the top 10 differential expressed genes per cell type. (E-G) Decidualization and decidualization rate after induction cell transplantation. Scale bar, 1 cm.

We further investigated the expression of critical transcription factors involved in trophoblast development within day 4.5 ci-MOs TE. In agreement with this we found that *Ascl2* (Bogutz et al., 2018), *Mbnl3* (Spruce et al., 2022), *Cdkn1c* were expressed in EPC clusters, especially in the migratory EPC; *Sox11*, *Igfbp2* (Jiang et al., 2023), *Fn1*(Lopez-Abad et al., 2016) were reported to be important to decidualization; and *Tfap2c*, *Sox21*, *Sox2* (Moretto Zita et al., 2015) were robustly expressed in TSC lineages (**Figure 6D**). At E7.5, *Sox21* RNA was highly expressed in the chorion but undetected in the ectoplacental cone (Moretto Zita et al., 2015). These results show that day 4.5 ci-MOs display trophoblast lineage development and the presence of distinct clusters, mirroring mouse placentation.

Although surrounded by PrE-like cells, we transplanted day 0 ci-MOs into the uterus of pseudo-pregnancy females as mural TE cells and EPC cells are found in day 0 and day 4.5 ci-MOs respectively. As expected, ci-MOs could not implant, likely due to lack of exposed mural TE and envelopment of ci-MOs by visceral endoderm. Nonetheless, efficient, up to 70%, and robust decidualization was observed, suggesting that ci-MOs may produce a currently unknown molecular stimulation for decidualization to occur (**Figure 6E-6G**). This phenotype aligns perfectly with our unexpected discovery of a distinct cluster of cells in day 4.5 ci-MOs co-expressing *Sox11*, *Igfbp2,* and *Fn1* representing decidualization-driving cells (**Figure S5D**). These cells were previously assumed to be associated with maternal cell contamination (Jiang et al., 2023) and we now propose that they may be produced by the embryo itself as it is present in our embryo model.

Together, these results show that ci-MOs have robust trophoblast lineage development.

### ci-MO developmental tree reconstruction

During embryo development, the zygote divides into a mass of cells from which the three blastocyst fates arise. These cells then develop into other types with distinct functions and molecular characteristics. Likewise, our chemical-induced ESCs generate progenitor cells capable of specifying the three blastocyst fates and subsequently self-assemble into an integrated embryo model closely resembling natural embryo development. To trace the transcriptional trajectories and dynamics underpinning ESCs till day 4.5 ci-MO development, we used a simulated diffusion-based computational approach, URD (Farrell et al., 2018). We collected and generated single-cell transcriptomes from 26818 cells, spanning 7 stages from the onset of uninduced ESC through D4.5 ci-MOs resembling late streak embryos (uninduced ESC; 24 h, 60 h, 96 h chemical induced cells; D1.5, D2.5 and D4.5 ci-MOs) (**Figure 7A**). This allowed us to reconstruct a developmental branching tree describing the transcriptional trajectories that lead to 24 distinct cell types across 16 branch points from uninduced cells through D4.5 gastrulating ci-MOs at 2-dimension and 3-dimension form (**Figure 7A and 7B**).

**Figure 7.**
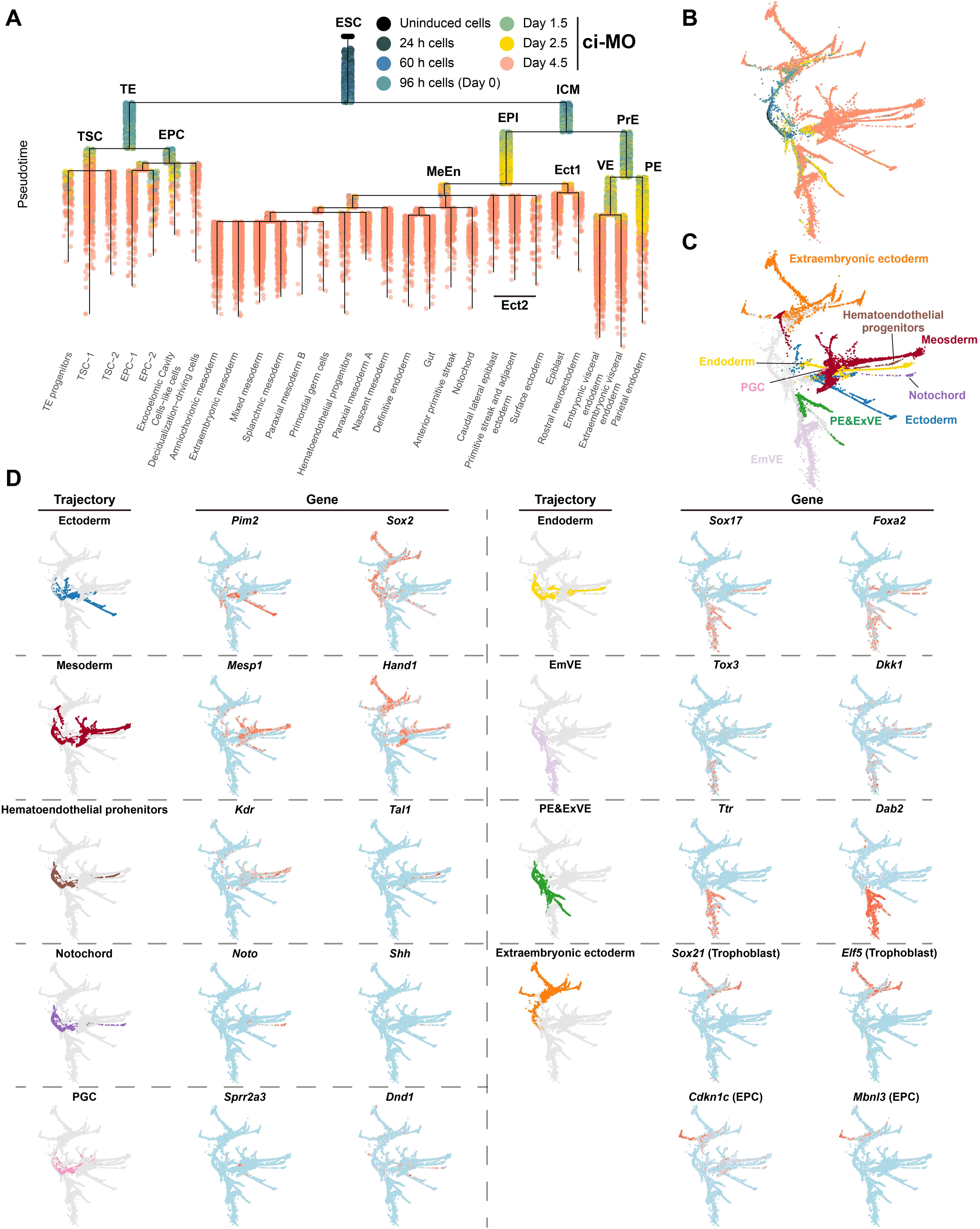
Reconstruction of transcriptional trajectories from chemically induced ESCs to day 4.5 ci-MOs. (A) URD differentiation tree for samples from uninduced cells to day 4.5 ci-MOs. (Uninduced cells, black; 24 h cells, dark slate grey; 60 h cells, steel blue; 96 h cells (D0), cadet blue; D1.5, dark sea blue; D2.5, gold; D4.5, Indian red; TE, trophoectoderm; ICM, inner cell mass; EPI, epiblast; PrE, Primitive endoderm; MeEn, Mesendoderm; Ect1/2, Ectoderm 1/2; VE, visceral endoderm; PE, Parietal endoderm.) (B) Force-directed layout of the URD differentiation tree, optimized for 2D visualization. (Uninduced cells, black; 24 h cells, dark slate grey; 60 h cells, steel blue; 96 h cells (D0), cadet blue; D1.5, dark sea blue; D2.5, gold; D4.5, Indian red.) (C) Force-directed layout of the URD differentiation tree, optimized for 2D visualization. (Ectoderm, blue; Mesoderm, red; Hematoendothelial progenitors, brown; Notochord, purple; PGC, pink; Endoderm, gold; Embryonic visceral endoderm (EmVE), light pink; Extraembryonic visceral endoderm (PE & ExVE), green; Extraembryonic ectoderm, orange.) (D) Trajectory of a specific population (or population group) identified by URD from roots (uninduced cells) to day 4.5 ci-MOs and gene expression of classical markers of that population. *Pim2* and *Sox2* mark Ectoderm, *Mesp1* and *Hand1* mark Mesoderm, *Kdr* and *Tal1* mark Hematoendothelial progenitors, *Noto* and *Shh* mark Notochord, *Sprr2a3* and *Dnd1* mark PGC, *Sox17* and *Foxa2* mark Endoderm, *Tox3* and *Dkk1* mark Embryonic visceral endoderm (EmVE), *Ttr* and *Dab2* mark Parietal endoderm and Extraembryonic visceral endoderm (PE & ExVE), *Sox21*, *Elf5* mark Trophoblast, *Cdkn1c* and *Mbnl3* mark EPC.

Notably, the first lineage branching occurred during the first pulse of chemical induction (60 h), segregating into TE-like and ICM-like lineages, which followed a similar route to the first lineage specification in vivo. Similarly, during the 96 hour chemical induction, a second branching in the ICM occurred, leading to the emergence of the embryo proper (EPI) and yolk sac lineages (PrE) (**Figure 7A**). Furthermore, in the EPI branches, the segregation into ectoderm 1 and mesendoderm was first observed; and the mesoderm, ectoderm 2, endoderm, and PGC were recovered in the embryo proper branch (**Figure 7A**). Obviously, the 3-dimension form also recapitulated the lineage segregation at gastrulating embryos with 3 germ layers, hematoendohelial progenitors, notochord, PGC, extraembryonic ectoderm, and extra-embryonic endoderm (**Figure 7C**).

We also used classic marker genes to confirm the annotations of the expected developmental trajectories of the tree (**Figure 7D**). For example, consistent with its known expression, the cardiac mesoderm marker gene *Hand1*(Barnes et al., 2010) was restricted primarily to two trajectories; the entire extraembryonic ectoderm and the later stages of mesoderm. In the TE branch, the developmental trajectory mirrored the early pathway of placental development, comprising not only the TSC derivatives, EPC but also EPC migratory derivatives (**Figure 6 and 7A**).

Meanwhile, in the PrE branch, the parietal endoderm is initially segregated, followed by the segregation of the visceral endoderm (**Figure 7A**). PrE and its derivatives are generally referred to as extra-embryonic endoderm, which contributes to extraembryonic lineages encompassing the visceral endoderm (VE) and the parietal endoderm (PE) (Morris et al., 2012; Thowfeequ et al., 2024). In a gastrulating embryo, the definitive endoderm (DE) forms by ingression of cells from the epiblast (Christodoulou et al., 2019). The DE comprises most of the cells of the gut and its accessory organs. Even though a large amount of overlap in markers between ExEn and DE, which makes it difficult to distinguish them, our developmental tree successfully tracked their origins and fates (Takaoka et al., 2011). This is also confirmed by the expression of ExVE and PE-specific markers (*Ttr* and *Dab2*), and endoderm marker *Sox17* and *Foxa2* in the respective trajectory branches. The tree further confirmed that AVE (marked by genes *Tox3* and *Dkk1* (Thowfeequ et al., 2024)) is not derived from distal VE but is formed from ExVE cells (**Figure 7D**).

We further identified the developmental trajectory of hematoendothelial progenitors and notochord, which developed from ICM-EPI-MeEn till the branching into mesoderm and endoderm by tracing the cells expressing the notochord marker gene *Noto(Beckers et al., 2007)* and the hematoendothelial progenitors makers *Kdr* (Li et al., 2017) and *Tal1* (Kallianpur et al., 1994) (**Figure 7D**).

Overall, our single-cell transcriptome analysis revealed that chemical induction, starting from ESCs, faithfully recapitulates the complex lineage branching and differentiation patterns observed in natural embryo development, underscoring the potential of ci-MOs to model embryogenesis at a single-cell resolution.

## Discussion

In this study, we explored the possibility of developing a high-fidelity embryo model that was transgene-free, cell mixing-free, direct, rapid and efficient, and easy to use in labs worldwide. We hypothesized that this could be achieved by chemically reprogramming ESCs into precursor-like cells of the three blastocyst fates. The expectation was that such cells could represent a more natural starting point and potentially better way of recreating embryo development in a petri dish. Strikingly, ci-MOs displayed unprecedented morphological and molecular resemblance to the late gastrulation embryo. This not only validated our hypothesis but also underscored the importance of the starting cell identity in determining the developmental potential of the embryo model.

ci-MOs were found to have a sophisticated TE compartment which featured, as in the embryo, an ectoplacental cone-like structure. Molecular analysis of this further confirmed the degree of TE development and sophistication. This data was unexpected as this structure in ci-MOs is completely enveloped by visceral endoderm and that way prevented from mimicking implantation. The interpretation we make is that TE development is independent of implantation and that the TE fate cells in our model are more developed than those in other embryo models. Robust and efficient decidualization was also detected upon the transfer of ci-MOs into the uteri of pseudo-pregnant mice indicating that potential signals prevenient from embryos to induce decidualization are also present in ci-MOs. Together, these results make ci-MOs a unique embryo model to study placenta development.

We focused our analysis on the late gastrulation stage. However, after additional culture, ci-MOs showed further developmental progression (data not included). In the future, it will be important to explore ci-MOs’ full potential by using improved apparatus and a wide range of additional culture conditions.

Similar to what happens in the embryo, ci-MOs PrE cells envelop the EPI and TE compartments. However, the embryo also forms an outer mural TE that it uses to attach to the uterus. Thus, despite our embryo model’s remarkable similarity to natural embryo development, it lacks a fundamental component necessary for implantation to occur. Thus, a strategy refinement to our model is needed to allow ci-MOs mural TE to adopt the correct position and thus enable development upon transfer into a uterus.

In conclusion, ci-MOs represent a high-fidelity embryo model with unique characteristics, and further refinement will help to unlock its full potential as an advanced model for broad biological research.

## Acknowledgments

We thank Naihe Jing for feedback on the manuscript. We thank Mengdan Zhang and Mingwei Min for their technical assistance. We thank the Advanced Bio-imaging Technology Platform and the flow cytometry core in Guangzhou National Laboratory for technical help. We appreciate the support from Xiongzhi Quan regarding mouse embryo microinjection and transplantation. This study was supported by grants from the Department of Science and Technology of Guangdong Province (2021ZT09Y233 to J.C.R.S), the National Natural Science Foundation of China (32100653 to H.L.), and Guangzhou National Laboratory (Major Project: GZNL2023A02005 & GZNL2023A02006, to J.C.R.S and H.L.). The J.C.R.S laboratory is supported by Guangzhou National Laboratory.

## Author contributions

Conceptualization, H.L. and J.C.R.S.; Methodology, H.L., J. H., W.G., J.W., and J.C.R.S.; Investigation, H.L., J. H., W.G., J.W, HP.L. and L.C.; Software, W.G. and J.W.; Writing – Original Draft, H.L. J. H., and J.C.R.S.; Writing – Review & Editing, H.L. and J.C.R.S.; Funding Acquisition, H.L. and J.C.R.S.; Resources, Y.G. and J.Z.; Supervision, H.L. and J.C.R.S.

## Figure legends

**Figure S1. Small molecule screen to identify compounds for inducing ESCs into blastocyst-like precursor cells. Related to Figure 1**.

(A) Different small molecules combination and immune-flow assay. K1, K2, K3, and K4 represent different combination and induction strategies, respectively. GATA6 is a primitive endoderm (PrE) marker, and CDX2 is a trophectoderm (TE) marker.
(B) Immunofluorescence of 60 h induction cells (OG2 ESCs). TE marker CDX2 (red); PrE marker GATA6 (white). ICM marker OCT4 (green) corresponds to the GFP signal driven by endogenous Oct4 activity; nuclear marker DAPI (cyan). Scale bar, 50 μm.
(C) Expression of cell lineage marker gene in K1, K2, K3, and K4 at different time points. Fold changes in mRNA expression of target genes were determined by ΔΔCt method using *Actin* as endogenous control.

(D-E) Immunofluorescence of 60 h induction cells (B6 and ICR ESCs). TE marker CDX2 (red); PrE marker GATA6 (white); ICM marker OCT4 (green); nuclear marker DAPI (cyan). Scale bar, 50 μm.

**Figure S2. Analysis of 96 h TE-like sub-clusters. Related to Figure 2**.

(A) UMAP of sub-clusters of 96 h TE-like cells.
(B) UMAP plots showing expression of Polar TE and mural TE markers in Sub-clusters of 96 h TE-like cells. *Cdx2*, *Ddah1*, *Gsto1*, *Ly6a* and *Ccnd1* mark Polar TE; *Krt18*, *Krt8*, *Itga6*, *Lgals1* and *Efnb1* mark mural TE.
(C) Selected enriched GO terms of sub-clusters of 96 h TE-like cells.
(D) Selected enriched KEGG terms of sub-clusters of 96 h TE-like cells.
(E) Immunofluorescence of 96 h induction cells. Mural TE marker KRT18 (yellow); polar TE marker CDX2 (purple); ICM marker OCT4 (green), nuclear marker DAPI (cyan). Scale bar, 50 μm.

**Figure S3. Immunofluorescence of ci-MOs at different stages. Related to Figure 4**.

(A-D) Immunofluorescence of day 1.5 and day 2.5 ci-MOs. AP2γ (red), OCT4 (green), GATA4 (white), CDX2 (red), GATA6 (white), DAPI (cyan). Scale bar, 100 μm.

(E) Immunofluorescence of day 3.5 ci-MOs. AP2γ (yellow), OCT4 (green), SOX2 (purple), and DAPI (cyan). Scale bar, 100 μm.

**Figure S4. Generation of ci-MOs in independent ESC cell lines. Related to Figure 4**.

(A-B) Immunofluorescence of day 1.5, day 2.5, and day 3.5 ci-MOs from K4.CDX2 (red), OCT4 (green), GATA4 (white), AP2γ (red), SOX2 (green), and DAPI (cyan). Scale bar, 100 μm.

## RESOURCE AVAILABILITY

### Lead contact

Further information and requests for resources or reagents should be directed to and will be fulfilled by the lead contact, José C. R. Silva.

### Materials availability

The mouse lines and other research reagents generated in this study will be distributed upon request to other research investigators under a Material Transfer Agreement.

### Data and code availability

The scRNA-seq data have been deposited in the Genome Sequence Archive in the National Genomics Data Center, China National Center for Bioinformation, Chinese Academy of Sciences (GSA: XXXXXXXXX). Any additional information required to reanalyze the data reported in this preprint is available from the corresponding authors upon request.

## EXPERIMENTAL MODEL AND STUDY PARTICIPANT DETAILS

### Animals

All animal experiments were performed following the Guide of the Care and Use of Laboratory Animals by the National Research Council and approved by the Institutional Animal Care and Use Committee of Guangzhou Institutes of Biomedicine and Health (GIBH), CAS. Any animals that appeared unhealthy before the start of experiments were excluded. No inclusion criterion was used. The ICR mice were purchased from Beijing Vital River Laboratory and the rats were purchased from Zhuhai Baishitong Biotechnology Co., Ltd. The mice were housed with a 12 h light/ dark cycle between 07:00 and 19:00 in a temperature-controlled room (22 ± 1 °C) with free access to water and food. CO_2_ inhalation was used for euthanasia. To ensure death following CO_2_ asphyxiation, cervical dislocation was performed.

### Cell lines

The following mouse ESC lines were used: The OG2 ESC line is a gift from Prof. Guangming Wu. OG2 ESCs were derived from 3.5 d.p.c ICM from 129 female mice crossing male OG2 mice.; Male ESC line), WT ICR ESC (ICR; Male ESC line) and WT B6 ESCs (C57B/6; Male ESC line). We note that in our hands, ESC lines with lower passages (within 20 passages) have much higher efficiency in yielding nice ci-Mos. E14tg2a ESC line responds well to our chemical cocktail induction, however, E14tg2a ESC line with 35 passages could not generate normal ci-MOs beyond day 5. All cell lines were routinely tested every two weeks to ensure that they were not contaminated with mycoplasma.

### H2B-mcherry OG2 cell line

CSII-EF-H2B-mcherry lentivirus were kindly provided by Mingwei Min’s lab. Fresh harvested H2B-mcherry lentivirus and N2B27 medium were mixed with 8 μg/mL polybrene (Biosharp, BL628A) and applied to dissociated OG2 ATV28 mESCs in suspension for 12 h. The supernatant was removed afterwards and refreshed with 2iLIF medium for 3 days. Then cells were purified by sorting and used for a chimera experiment.

## Methods details

### Mouse ESC culture conditions

All mouse ESCs were cultured in 2iLIF medium on gelatin-coated plates in N2B27 basal medium supplemented with 20 ng/mL murine LIF (qKine, Qk018), 3 μM CHIR99021 (Selleck, S2924) and 1 μM PD0325901 (Selleck, S1036). N2B27 basal medium comprised a 1:1 mix of DMEM/F12 (Gibco, C11330500CP) and Neurobasal A (Gibco, 21103049) supplemented with 1×B27 (Gibco, 17504044), 1×N2 (Gibco, 17502048), 1×GlutaMAX (Gibco, 35050061), 1×MEM NEAA (Gibco, 11140050), 0.1 mM 2-mercaptoethanol (Gibco, 21985023). Cells were refreshed medium daily and cultured under 20% O_2_ and 5% CO_2_ at 37 °C.

### Chemical induction of mouse ESC

For chemical induction, ESCs were dissociated using accutase as previously described (Li et al., 2024). Dissociated ESCs were seeded at a density of 2.0×10^5^ per well on gelatin-coated 6-well plates in chemical cocktails. Small molecules were added to the N2B27 Basal medium. Cells were used for further experiments after 60 h induction. The details of each cocktail were listed.

CL6A medium: N2B27 supplemented with 3 μM CHIR99021 (Selleck, S2924), 20 ng/mL murine LIF (qKine, Qk018), 10 μM E616452 (Selleck, S7223) and 0.05 μM AM580 (Selleck, S2933). C6A medium: N2B27 supplemented with 3 μM CHIR99021, 10 μM E616452 and 0.05 μM AM580. C^+^6A medium : supplemented with 6 μM CHIR99021, 10 μM E616452 and 0.05 μM AM580. C^++^6A medium : N2B27 supplemented with 9 μM CHIR99021, 10 μM E616452, and 0.05 μM AM580.C^+^L6A medium : N2B27 supplemented with 6 μM CHIR99021, 20 ng/mL murine LIF, 10 μM E616452 and 0.05 μM AM580.

K1 treatment: induce the ESCs with CL6A medium for 60 h; K2 treatment: induce the cells with C6A medium for 24 h, followed with 36 h treatment with CL6A; K3 treatment: induce the cells with C^+^6A medium for 24 h, followed with 36 h treatment with CL6A; K4 treatment: induce the cells with C^++^6A medium for 24 h, followed with 36 h treatment with CL6A; K5 treatment: induce the cells with C^+^6A medium for 24 h, followed with 36 h treatment with C^+^L6A.

The 60 h chemical-induced cells were dissociated into single-cell suspension filtered through 40 μm strainers. Cells were cultured at a density of 3-3.5×10^4^ per Aggrewell with PM medium at 37 °C, 5% CO2 culture condition for 1.5 days. FC medium: Dulbecco’s modified essential medium (Gibco 41966052) supplemented with 20% (v/v) FBS (Carpicorn scientific, FBS-52A), 1 mM sodium pyruvate (Gibco, 11360039), 2 mM GlutaMAX (Gibco, 35050038), 1% MEM non-essential amino acids (Gibco, 11140035), 0.1 mM 2-mercaptoethanol (Gibco, 31350010) and 1% (v/v) penicillin/streptomycin (Gibco, 15140122).

PM medium: FC medium supplemented with 25 ng/mL rhFGF4 (R&D, 235-F4), 10 ng/mL hBMP4 (R&D, 314-BP-010/CF), 20 ng/mL Activin A (PeproTech, 120-14E), 3 μM XAV939 (Sigma, X3004), 2 μM TRULI (Selleck, E1061), 1 μg/mL Heparin (Sigma, H3149), 200 μM L-ascorbic acid 2-phosphate (Sigma, A8960) and 1 × ITS-G (Gibco, 41400045).

### In-house rat serum preparation

Rat serum was prepared with reference to the methods of Takahashi *et al* (Takahashi et al., 2014). The specific methods were as follows. After rats were anesthetized with 2.5%-4% isofluorane, abdominal aortic blood was collected using a 0.5 mm venous blood collection needle (SANLI, F0326-9-1) and a coagulant tube (BD Biosciences, 367955). After each tube was collected and mixed up 6-7 times, it was allowed to stand at room temperature for at least 30 min. When it has obvious stratification, centrifuge the test tubes for 15 min at 2,000 × *g* at room temperature and then carefully transfer the serum into a 15 mL sterile tube (Corning^®^, 430053) using a sterile pipette in a laminar air-flow cabinet. Incubate the serum in a water bath at 55 °C for 45 min to inactivate the complement system. Finally, inactivated rat serum was frozen in liquid nitrogen and stored at -80 °C, defrosted before use, and filtered with 0.22 μm filters. Specific pathogen-free, 9-week-old (>300 *g*) male rats were used in this study.

### Chemical induction-derived integrated embryo model (ci-MO) culture in vitro

To generated a chemical induction-derived integrated embryo model, the protocol was modified and adapted from the previously described (Lau et al., 2022; Li et al., 2023). At day 0, wash the day 0 ciMO twice with FC medium and fresh 1 mL FC medium to each well.

At day 1.5, fresh 1.5 mL IVC1 medium to each well.

At day 2.5, the ci-MOs were transferred to a suspension 6-well plate, fresh 4 mL IVC1 medium each well, and cultured in a shaking incubator at 80 rpm, 37°C and 5% CO_2_ condition.

From day 3.5, ci-MOs were transferred to a 5 mL bioreactor (ABLE Biott, BWV-S005A), fresh 3 mL IVC2 medium each well and cultured in a Bioreactor Magnetic Stir System-6ch. Base plate for 5 mL (ABLE Biott, BWS-S03N0S-6B) at 50 rpm, 37°C, and 5% CO_2_ condition.

Day 1.5, day 2.5, day 3.5, and day 4.5 ci-MOs were collected for further experiments and the details of each medium were as follows.

IVC1 medium: Advanced DMEM/F12 (Gibco, 12634028) supplemented with 20% (v/v) FBS (Carpicorn scientific, FBS-52A), 2 mM GlutaMax (Gibco 35050038), 1 × ITS-X (Gibco, 51500056), 100 nM T3 (3,3′,5-Triiodo-L-thyronine sodium salt, sigma, T6397), 8 nM β-estradiol (Sigma, E8875), 200 ng/mL progesterone (Sigma, P0130) and 25 μM N-acetyl-L-cysteine (Sigma, C7880), 1 mg/mL D-glucose (Sigma, G8644) and 1% (v/v) penicillin–streptomycin (Gibco, 15140122).

IVC2 medium: DMEM (low glucose, pyruvate, no glutamine, no phenol red, Gibco, 11880) supplemented with 50% rat serum, 30% human serum AB (Off the Clot, Gemini, 100-318), 2 mM GlutaMax (Gibco 35050038), 1 mM sodium pyruvate (Gibco 11360039), 11 mM HEPES (Gibco, 15630130), 4 mg/mL D-glucose (sigma, G8644) and 1% (v/v) penicillin–streptomycin (Gibco, 15140122).

### Immunostaining of cells and whole-mount immunostaining of ci-MOs

Cells were cultured in 96-well plate and fixed with 4% PFA for 30 min at room temperature, while ci-MOs were fixed with 4% PFA overnight at 4 °C. Then samples were blocked with blocking buffer (DPBS containing 5% FBS, 5% Normal Goat Serum, 2% BSA, 0.3% Triton X-100) at room temperature for 1 hour, followed by incubating with primary antibodies diluted in blocking buffer at 4 °C overnight. Samples were washed three times with PBST (DPBS containing 0.05% Tween-20), then were incubated with Alexa Fluor tagged secondary antibodies against the appropriated species at 1:1000 for 1 h at room temperature. Samples were washed three times with PBST after samples were stained with DAPI. For natural embryos up to E6.5 and ci-MOs up to day 5, sample were treated with an extra clearing step: gradient treatment with 25%, 50%, 75%, 95%, and 100% 2, 2′-Thiodiethanol (v/v, diluted in DPBS; sigma, 166782). Samples were imaged on confocal microscopes FV3000 (Olympus) and Image J software (NIH, USA) was used to analyze. The details of antibodies and their dilutions are provided in Key Resources Table.

### Immunohistochemistry for ci-MOs paraffin section

ci-MOs of various developmental stages were fixed in 4% PFA (Beyotime, P0099) for overnight at 4 °C. ci-MOs embedded in paraffin were sectioned into 4 μm slices with a microtome (Leica, Germany) and processed for deparaffinization and rehydration followed by antigen retrieval in 10 mM sodium citrate (pH 6.0, Servicebio Biological, G1202-250ML) (Buttler et al., 2008). Cooled and washed three times with phosphate buffer (PBS) (pH 7.4), 5 minutes each time. The following steps are same with previous methods (Immunostaining of cells and whole-mount immunostaining of ci-MOs) until the secondary antibody and DAPI incubated. Then cover the slips with an anti-fade mounting medium. All sections were imaged on confocal microscopes EVIDENT FV3000 and Image J software (NIH, USA) was used to analyze. The details of antibodies and their dilutions are provided in Key Resources Table.

### Eight-cell injection and chimeric embryo generation

Mouse 8-cell embryos were flushed from uterus/oviducts of plugged CD-1 (ICR) mice using M2 medium (Millipore, MR-015-D) and cultured in KSOM-AA (Millipore, MR-106-D) covered by mineral oil (Sigma, M8410) at 37 °C and 5% CO_2_. Microinjections were performed using a FemtoJet (Eppendorf).

To obtain chimeric blastocysts, four to six healthy cells were injected into each 8-cell stage embryo perivitelline space. The injected embryos were cultured in KSOM-AA drops for an additional 60 h. To obtain E6.5-7.5 chimera, after injection, the embryos were removed to micro drops of KSOM-AA culture medium and placed in a humidified incubator with 5% CO_2_ at 37 °C. After a 2 h of incubation, about 15 chimeric embryos were transferred to each surrogate mouse by uterine transfer, and the conceptuses were dissected at E6.5/ 7.5 developmental stages.

### Flow cytometry and FACS sorting

Cells were dissociated with accutase and suspended in appropriate medium. Cell immune flow was blocked with blocking buffer (5% FBS, 5% Normal Goat Serum, 2% BSA in DPBS without Ca^2+^ and Mg^2+^) for 20min, then primary antibody diluted with blocking buffer was added and incubated for 1 h at room temperature. After washing three times with DPBS without Ca^2+^ and Mg^2+^, the secondary antibody diluted with blocking solution was added and incubated for 30 min. Washed three times with DPBS without Ca^2+^ and Mg^2+^ and cells filtered through 40 μm strainers. Samples were analyzed by BD LSRFortessa X-20 and data analysis was performed via FlowJo software. For FACS sorting, single cell suspension was analyzed by BD FACSAria and target cell population was sorted for subsequent experiment.

### RNA extraction & RT-qPCR analysis

Total RNA was extracted using Ultrapure RNA Kit (CWBIO, CW0581M). 1 μg of total RNA was reverse transcribed into cDNA using RevertAid™ First Strand cDNA Synthesis Kit (Thermo, K16225) according to the manufacturer’s instructions. qRT-PCR was performed using TB Green^®^ Premix Ex Taq™ II (Takara, RR820A) and QuantStudio 3 Real-Time PCR System (Applied Biosystems). Fold change in mRNA expression of target genes were determined by ΔΔCt method using *Actin* as endogenous control. qPCR primers are listed in **Table 1**.

**Table 1.**
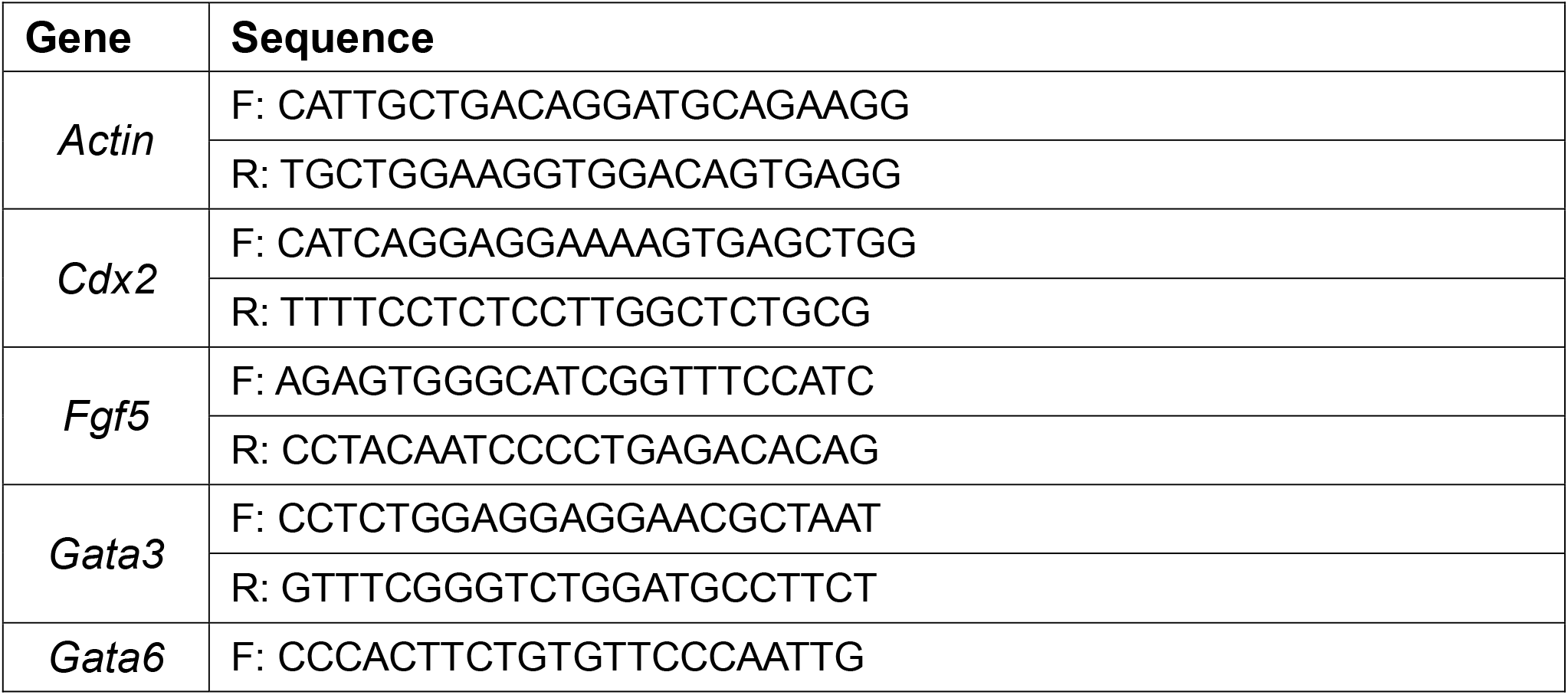
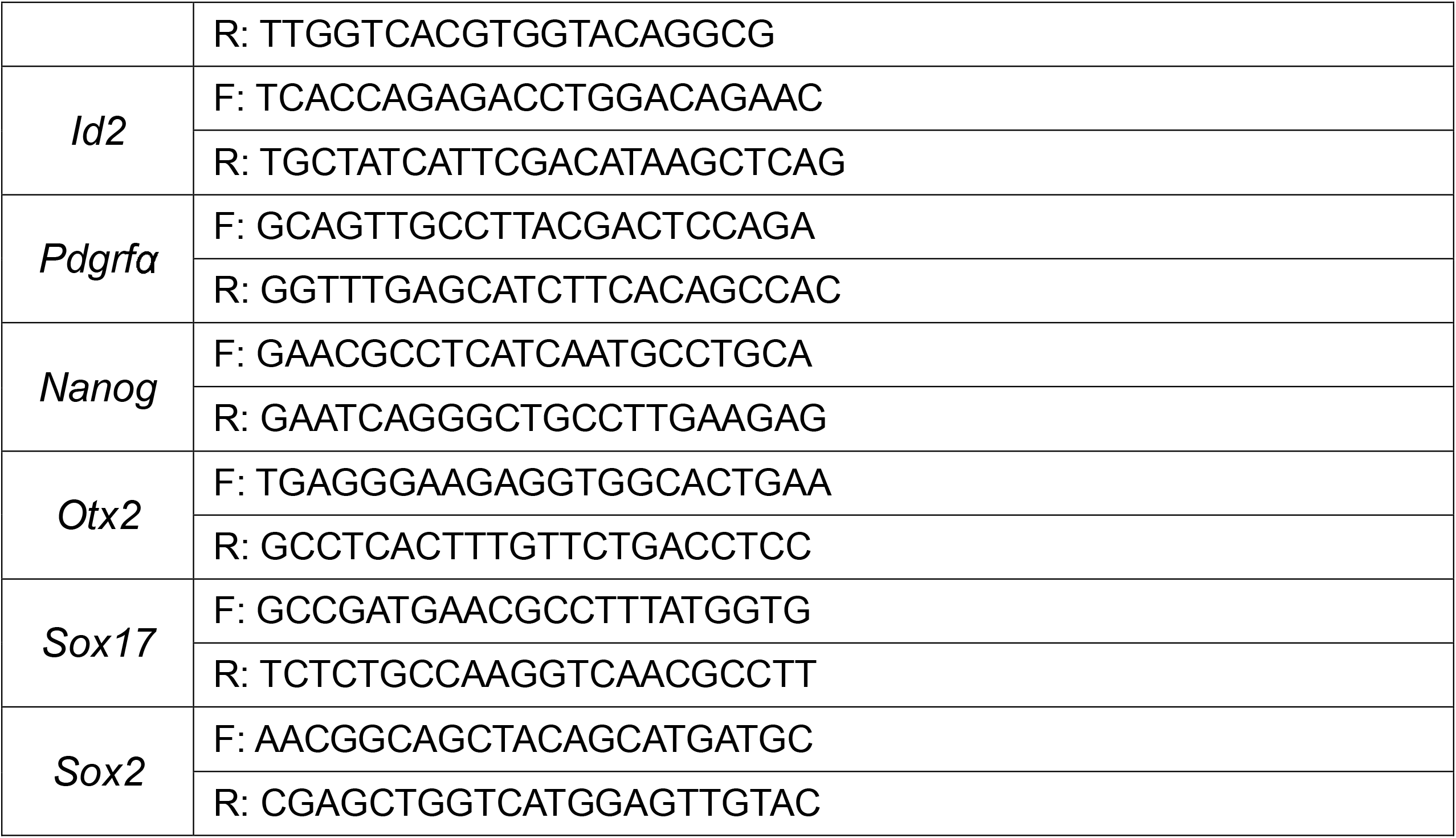
List of qPCR primers.

### Preparation of single cell suspensions

For chemical induced cells, cells were dissociated with accutase. For ciMOs, ciMOs were digested with 100 μL 10 U/ml papain (Worthington Biochemicals, LS003126) at 37 °C for 15-30 min.

Single cells were washed twice with cold PBS, filtered through a 35 μm cell strainer and loaded onto 10× Genomics Chromium system using Single Cell 3’ Reagent Kits V3.1 at a concentration of 16,000 single cells per sample. Libraries were sequenced on MGISEQ2000 (MGI Tech) system in PE100 mode.

### Pre-processing Single cell RNA-seq data

#### Single-cell RNA-seq data generated in this study

All single-cell RNA-seq data in this study were analyzed using a custom reference genome built in accordance with the instructions from 10× Genomics website (https://support.10xgenomics.com/single-cell-gene-expression/software/pipelines/latest/using/tutorial_mr). Pre-built genome sequence and gene annotations were downloaded from Ensembl GRCm39 release 104. Raw FASTQ data were quality controlled using fastp(Chen et al., 2018) (v.0.21.0, https://github.com/OpenGene/fastp) with default parameters. Alignment, annotation, PCR duplicate removal and gene expression quantification were subsequently performed using the STARsolo pipeline (https://github.com/alexdobin/STAR/blob/master/docs/STARsolo.md) with STAR (Dobin et al., 2013) (v.2.7.9a).

#### Public Single-cell RNA-seq data

Public data from GSE45719 (Deng et al., 2014), GSE84892 (Posfai et al., 2017), GSE109071 (Cheng et al., 2019), GSE100597 (Mohammed et al., 2017), GSE123046 (Nowotschin et al., 2019), E-MTAB-6967 (Pijuan-Sala et al., 2019) (mouse embryo stages ranging from zygote to E7.5 with all known lineages within this period) were downloaded and pre-processed with the same protocol as described for data generated in this study or with adaptation following the authors instructions.

### Downstream analysis of scRNA-seq data

#### Linear dimensional reduction with Principal Component Analysis (PCA)

Further analysis was performed in R 4.2.2 with Seurat (Hao et al., 2021) (v.4.3.0). Based on initial evaluation of per-cell quality control metrics, cells with less than 4000 (Smart-seq2 data) or 2000 (10× Genomics data) detected genes or >= 5% mitochondrial gene percentage were filtered out. Genes detected in at least 3 cells were retained. Log-normalization, centering and scaling of the filtered count data were performed with NormalizeData and ScaleData functions sequentially and top 2000 highly variable genes were selected with FindVariableFeatures function. Principal component analysis (PCA) was applied on the scaled data using RunPCA function in Seurat with default parameters based on the selected highly variable genes. The PCA coordinates were later used as input of graph-based clustering approach.

#### Batch effect correction and integration of Smart-seq2 and 10× Genomics experiments from multiple sources

For comparisons between day 4.5 ci-MOs and public natural embryo datasets, canonical correlation analysis (CCA) approach, standard integration pipelines described in Seurat official website (https://satijalab.org/seurat/articles/integration_introduction.html) were used for anchors determination and data-sets alignment with default parameters. For the integration of public data covering different lineages and developmental stages from multiple sources as was previously described, fastMNN (Haghverdi et al., 2018) approach was employed. Each dataset from different experiments was considered a batch and contained at least one shared cell population with the others. Log-normalization was performed using computeSumFactors function from scran package (Lun et al., 2016) (v.1.20.1) and scaling normalization across batches was then performed with multiBatchNorm function in batchelor package(Haghverdi et al., 2018) (v.1.8.1). The log-normalized, batch-effect corrected datasets were thereafter integrated using the fastMNN approach implemented via SeuratWrappers (v.0.3.0, https://github.com/satijalab/seurat-wrappers). The MNN low-dimensional coordinates were later used for graph-based clustering and visualization.

#### Graph-based clustering and visualization with Uniform Manifold Approximation and Projection (UMAP)

Shared nearest neighbor (SNN) graphs were constructed based on the dimensionally reduced and batch effect corrected data with FindNeighbors function and were subsequently partitioned via Louvain algorithm implemented with FindClusters function at a range of resolutions selected according to library size and data heterogeneity. UMAP was implemented via the RunUMAP function for visualization.

#### Lineage trajectory analysis

For developmental trajectories inference of ci-MOs, URD (Farrell et al., 2018) (v.1.1.1, https://github.com/farrellja/URD) was chosen. To perform URD analysis, we followed the detailed instructions provided by the author on URD from GitHub (https://github.com/farrellja/URD/blob/master/Analyses/QuickStart/URD-QuickStart-AxialMesoderm.md). We assigned a group of naïve stem cells with the signature of Cdx2^-^/Gata6^-^/Pou5f1^+^/Nanog^+^ as ‘uninduced cells’ as root, and day 4.5 ci-MOs as tips, resulting in a branching trajectory tree. For RNA velocity calculation, we used scvelo (Bergen et al., 2020) (v.0.3.1, https://github.com/theislab/scvelo), following the author’s instruction on https://scvelo.readthedocs.io/en/stable/index.html.

#### Lineage score calculation

To measure the similarity of our 24 h, 60 h, and 96 h cells with natural embryos, we used the top 50 differentially expressed genes of E3.5 and E4.5 mouse embryos to calculate the lineage scores of each single cell. In detail, the lineage scores were calculated using the AddModuleScore function of the Seurat package in each cell for these lineage-specific genes which were then normalized between 1 and 100 with E4.5 blastocysts cells as maximum (Figure 2F) or within our samples across induction time (Figure S2A). Next, the ‘ggtern’ package (v3.5.0) was used to visualize and transform the three-dimensional axis into two dimensions. Each dot represents one cell and colored with its lineage identity (Figure 2F).

## QUANTIFICATION AND STATISTICAL ANALYSIS

All statistic results and graphs were generated by Excel or R. The numbers of samples and types of statistical analyses are given in figure legends and results sections.

## Supplemental information

**Table S1**. Conserved genes between E4.5 blastocysts and 96 h cells, related to Figure 2H.

**Table S2**. Top differentially expressed genes of E3.5 and E4.5 mouse embryos, related to Figure 3E.

**Table S3**. Top GO and KEGG terms of subclusters of 96 h TE-like cells, related to Figure S2C and S2D.

**Table S4**. Top 30 differential expressed gene list of day 4.5 ci-MOs (OG2 ESC), related to Figure 5D.

**Table S5**. Top GO terms of day 4.5 ci-MOs (OG2 ESC), related to Figure 5D.

**Table S6**. Top 30 differential expressed gene list of Day 4.5 ci-MOs (B6 ESC), related to Figure S5D.

**Table S7**. Top GO terms of day 4.5 ci-MOs (B6 ESC), related to Figure S5D.

## References

Ajima, R., Sakakibara, Y., Sakurai-Yamatani, N., Muraoka, M., and Saga, Y. (2021). Formal proof of the requirement of MESP1 and MESP2 in mesoderm specification and their transcriptional control via specific enhancers in mice. Development 148.

Amadei, G., Handford, C.E., Qiu, C., De Jonghe, J., Greenfeld, H., Tran, M., Martin, B.K., Chen, D.Y., Aguilera-Castrejon, A., Hanna, J.H., et al. (2022). Embryo model completes gastrulation to neurulation and organogenesis. Nature 610, 143–153.

Amadei, G., Lau, K.Y.C., De Jonghe, J., Gantner, C.W., Sozen, B., Chan, C., Zhu, M., Kyprianou, C., Hollfelder, F., and Zernicka-Goetz, M. (2021). Inducible Stem-Cell-Derived Embryos Capture Mouse Morphogenetic Events In Vitro. Dev Cell 56, 366–382 e369.

Arnold, S.J., and Robertson, E.J. (2009). Making a commitment: cell lineage allocation and axis patterning in the early mouse embryo. Nat Rev Mol Cell Biol 10, 91–103.

Artus, J., Douvaras, P., Piliszek, A., Isern, J., Baron, M.H., and Hadjantonakis, A.K. (2012). BMP4 signaling directs primitive endoderm-derived XEN cells to an extraembryonic visceral endoderm identity. Dev Biol 361, 245–262.

Bardot, E.S., and Hadjantonakis, A.K. (2020). Mouse gastrulation: Coordination of tissue patterning, specification and diversification of cell fate. Mech Dev 163, 103617.

Barnes, R.M., Firulli, B.A., Conway, S.J., Vincentz, J.W., and Firulli, A.B. (2010). Analysis of the Hand1 cell lineage reveals novel contributions to cardiovascular, neural crest, extra-embryonic, and lateral mesoderm derivatives. Dev Dyn 239, 3086–3097.

Beccari, L., Moris, N., Girgin, M., Turner, D.A., Baillie-Johnson, P., Cossy, A.C., Lutolf, M.P., Duboule, D., and Arias, A.M. (2018). Multi-axial self-organization properties of mouse embryonic stem cells into gastruloids. Nature 562, 272–276.

Beckers, A., Alten, L., Viebahn, C., Andre, P., and Gossler, A. (2007). The mouse homeobox gene Noto regulates node morphogenesis, notochordal ciliogenesis, and left right patterning. Proc Natl Acad Sci U S A 104, 15765–15770.

Belo, J.A., Bouwmeester, T., Leyns, L., Kertesz, N., Gallo, M., Follettie, M., and De Robertis, E.M. (1997). Cerberus-like is a secreted factor with neutralizing activity expressed in the anterior primitive endoderm of the mouse gastrula. Mech Dev 68, 45–57.

Bergen, V., Lange, M., Peidli, S., Wolf, F.A., and Theis, F.J. (2020). Generalizing RNA velocity to transient cell states through dynamical modeling. Nat Biotechnol 38, 1408–1414.

Bogutz, A.B., Oh-McGinnis, R., Jacob, K.J., Ho-Lau, R., Gu, T., Gertsenstein, M., Nagy, A., and Lefebvre, L. (2018). Transcription factor ASCL2 is required for development of the glycogen trophoblast cell lineage. PLoS Genet 14, e1007587.

Boroviak, T., Stirparo, G.G., Dietmann, S., Hernando-Herraez, I., Mohammed, H., Reik, W., Smith, A., Sasaki, E., Nichols, J., and Bertone, P. (2018). Single cell transcriptome analysis of human, marmoset and mouse embryos reveals common and divergent features of preimplantation development. Development 145.

Buttler, K., Ezaki, T., and Wilting, J. (2008). Proliferating mesodermal cells in murine embryos exhibiting macrophage and lymphendothelial characteristics. BMC Dev Biol 8, 43.

Cao, N., Huang, Y., Zheng, J., Spencer, C.I., Zhang, Y., Fu, J.D., Nie, B., Xie, M., Zhang, M., Wang, H., et al. (2016). Conversion of human fibroblasts into functional cardiomyocytes by small molecules. Science 352, 1216–1220.

Chen, S., Zhou, Y., Chen, Y., and Gu, J. (2018). fastp: an ultra-fast all-in-one FASTQ preprocessor. Bioinformatics 34, i884–i890.

Cheng, L., Hu, W., Qiu, B., Zhao, J., Yu, Y., Guan, W., Wang, M., Yang, W., and Pei, G. (2014). Generation of neural progenitor cells by chemical cocktails and hypoxia. Cell Res 24, 665–679.

Cheng, S., Pei, Y., He, L., Peng, G., Reinius, B., Tam, P.P.L., Jing, N., and Deng, Q. (2019). Single-Cell RNA-Seq Reveals Cellular Heterogeneity of Pluripotency Transition and X Chromosome Dynamics during Early Mouse Development. Cell Rep 26, 2593–2607 e2593.

Christodoulou, N., Weberling, A., Strathdee, D., Anderson, K.I., Timpson, P., and Zernicka-Goetz, M. (2019). Morphogenesis of extra-embryonic tissues directs the remodelling of the mouse embryo at implantation. Nat Commun 10, 3557.

Deng, Q., Ramskold, D., Reinius, B., and Sandberg, R. (2014). Single-cell RNA-seq reveals dynamic, random monoallelic gene expression in mammalian cells. Science 343, 193–196.

Dobin, A., Davis, C.A., Schlesinger, F., Drenkow, J., Zaleski, C., Jha, S., Batut, P., Chaisson, M., and Gingeras, T.R. (2013). STAR: ultrafast universal RNA-seq aligner. Bioinformatics 29, 15–21.

Farrell, J.A., Wang, Y., Riesenfeld, S.J., Shekhar, K., Regev, A., and Schier, A.F. (2018). Single-cell reconstruction of developmental trajectories during zebrafish embryogenesis. Science 360.

Fu, Y., Huang, C., Xu, X., Gu, H., Ye, Y., Jiang, C., Qiu, Z., and Xie, X. (2015). Direct reprogramming of mouse fibroblasts into cardiomyocytes with chemical cocktails. Cell Res 25, 1013–1024.

Gao, Y., Han, W., Dong, R., Wei, S., Chen, L., Gu, Z., Liu, Y., Guo, W., and Yan, F. (2024). Efficient Reprogramming of Mouse Embryonic Stem Cells into Trophoblast Stem-like Cells via Lats Kinase Inhibition. Biology (Basel) 13.

Guo, M., Wu, J., Chen, C., Wang, X., Gong, A., Guan, W., Karvas, R.M., Wang, K., Min, M., Wang, Y., et al. (2024). Self-renewing human naive pluripotent stem cells dedifferentiate in 3D culture and form blastoids spontaneously. Nat Commun 15, 668.

Haghverdi, L., Lun, A.T.L., Morgan, M.D., and Marioni, J.C. (2018). Batch effects in single-cell RNA-sequencing data are corrected by matching mutual nearest neighbors. Nat Biotechnol 36, 421–427.

Hannibal, R.L., and Baker, J.C. (2016). Selective Amplification of the Genome Surrounding Key Placental Genes in Trophoblast Giant Cells. Curr Biol 26, 230–236.

Hao, Y., Hao, S., Andersen-Nissen, E., Mauck, W.M., 3rd, Zheng, S., Butler, A., Lee, M.J., Wilk, A.J., Darby, C., Zager, M., et al. (2021). Integrated analysis of multimodal single-cell data. Cell 184, 3573–3587 e3529.

Harrison, S.E., Sozen, B., Christodoulou, N., Kyprianou, C., and Zernicka-Goetz, M. (2017). Assembly of embryonic and extraembryonic stem cells to mimic embryogenesis in vitro. Science (New York, NY) 356, 6334 eaal1810.

Hayakawa, K., Himeno, E., Tanaka, S., and Kunath, T. (2015). Isolation and manipulation of mouse trophoblast stem cells. Curr Protoc Stem Cell Biol 32, 1E 4 1–1E 4 32.

Hou, P., Li, Y., Zhang, X., Liu, C., Guan, J., Li, H., Zhao, T., Ye, J., Yang, W., Liu, K., et al. (2013). Pluripotent stem cells induced from mouse somatic cells by small-molecule compounds. Science 341, 651–654.

Hu, Y., Yang, Y., Tan, P., Zhang, Y., Han, M., Yu, J., Zhang, X., Jia, Z., Wang, D., Yao, K., et al. (2023). Induction of mouse totipotent stem cells by a defined chemical cocktail. Nature 617, 792–797.

Iturbide, A., Ruiz Tejada Segura, M.L., Noll, C., Schorpp, K., Rothenaigner, I., Ruiz-Morales, E.R., Lubatti, G., Agami, A., Hadian, K., Scialdone, A., et al. (2021). Retinoic acid signaling is critical during the totipotency window in early mammalian development. Nat Struct Mol Biol 28, 521–532.

Jiang, X., An, W., Yang, X., Lin, J., Ma, S., Wang, D., and Tang, S. (2018). Asymmetric distribution of CRUMBS polarity complex proteins from compacted 8-cell to blastocyst stage during mouse preimplantation development. Gene Expr Patterns 27, 93–98.

Jiang, X., Wang, Y., Xiao, Z., Yan, L., Guo, S., Wang, Y., Wu, H., Zhao, X., Lu, X., and Wang, H. (2023). A differentiation roadmap of murine placentation at single-cell resolution. Cell Discov 9, 30.

Kallianpur, A.R., Jordan, J.E., and Brandt, S.J. (1994). The SCL/TAL-1 gene is expressed in progenitors of both the hematopoietic and vascular systems during embryogenesis. Blood 83, 1200–1208.

Kelly, G.M., Saijoh, Y., Finkielsztein, A., and Mangos, S. (2008). Mouse G-protein gamma3 expression in the developing CNS and neural crest cell derivatives. Int J Dev Biol 52, 1143–1150.

Kempf, H., Olmer, R., Haase, A., Franke, A., Bolesani, E., Schwanke, K., Robles-Diaz, D., Coffee, M., Gohring, G., Drager, G., et al. (2016). Bulk cell density and Wnt/TGFbeta signalling regulate mesendodermal patterning of human pluripotent stem cells. Nat Commun 7, 13602.

Klaffky, E., Williams, R., Yao, C.C., Ziober, B., Kramer, R., and Sutherland, A. (2001). Trophoblast-specific expression and function of the integrin alpha 7 subunit in the peri-implantation mouse embryo. Dev Biol 239, 161–175.

Kurimoto, K., Yabuta, Y., Ohinata, Y., Ono, Y., Uno, K.D., Yamada, R.G., Ueda, H.R., and Saitou, M. (2006). An improved single-cell cDNA amplification method for efficient high-density oligonucleotide microarray analysis. Nucleic Acids Res 34, e42.

Langkabel, J., Horne, A., Bonaguro, L., Holsten, L., Hesse, T., Knaus, A., Riedel, Y., Becker, M., Handler, K., Elmzzahi, T., et al. (2021). Induction of Rosette-to-Lumen stage embryoids using reprogramming paradigms in ESCs. Nat Commun 12, 7322.

Lau, K.Y.C., Rubinstein, H., Gantner, C.W., Hadas, R., Amadei, G., Stelzer, Y., and Zernicka-Goetz, M. (2022). Mouse embryo model derived exclusively from embryonic stem cells undergoes neurulation and heart development. Cell Stem Cell 29, 1445–1458 e1448.

Lee, M.B., Kooistra, M., Zhang, B., Slow, S., Fortier, A.L., Garrow, T.A., Lever, M., Trasler, J.M., and Baltz, J.M. (2012). Betaine homocysteine methyltransferase is active in the mouse blastocyst and promotes inner cell mass development. J Biol Chem 287, 33094–33103.

Lewis, S.L., Khoo, P.L., Andrea De Young, R., Bildsoe, H., Wakamiya, M., Behringer, R.R., Mukhopadhyay, M., Westphal, H., and Tam, P.P. (2007). Genetic interaction of Gsc and Dkk1 in head morphogenesis of the mouse. Mech Dev 124, 157–165.

Li, H., Chang, L., Huang, J., and Silva, J.C.R. (2024). Protocol for generating mouse morula-like cells resembling 8- to 16-cell stage embryo cells. STAR Protoc 5, 102934.

Li, H., Chang, L., Wu, J., Huang, J., Guan, W., Bates, L.E., Stuart, H.T., Guo, M., Zhang, P., Huang, B., et al. (2023). In vitro generation of mouse morula-like cells. Dev Cell 58, 2510–2527 e2517.

Li, Y., Brauer, P.M., Singh, J., Xhiku, S., Yoganathan, K., Zuniga-Pflucker, J.C., and Anderson, M.K. (2017). Targeted Disruption of TCF12 Reveals HEB as Essential in Human Mesodermal Specification and Hematopoiesis. Stem Cell Reports 9, 779–795.

Lindsley, R.C., Gill, J.G., Kyba, M., Murphy, T.L., and Murphy, K.M. (2006). Canonical Wnt signaling is required for development of embryonic stem cell-derived mesoderm. Development 133, 3787–3796.

Linneberg-Agerholm, M., Sell, A.C., Redo-Riveiro, A., Proks, M., Knudsen, T.E., Perera, M., and Brickman, J.M. (2023). Enhancer status in the primitive endoderm supports unrestricted lineage plasticity in regulative development. 2023.2005.2020.540779.

Litwin, M., Clark, K., Noack, L., Furze, J., Berndt, M., Albelda, S., Vadas, M., and Gamble, J. (1997). Novel cytokine-independent induction of endothelial adhesion molecules regulated by platelet/endothelial cell adhesion molecule (CD31). J Cell Biol 139, 219–228.

Lopez-Abad, M., Iglesias-Platas, I., and Monk, D. (2016). Epigenetic Characterization of CDKN1C in Placenta Samples from Non-syndromic Intrauterine Growth Restriction. Front Genet 7, 62.

Lun, A.T., McCarthy, D.J., and Marioni, J.C. (2016). A step-by-step workflow for low-level analysis of single-cell RNA-seq data with Bioconductor. F1000Res 5, 2122.

McClatchie, T., Meredith, M., Ouedraogo, M.O., Slow, S., Lever, M., Mann, M.R.W., Zeisel, S.H., Trasler, J.M., and Baltz, J.M. (2017). Betaine is accumulated via transient choline dehydrogenase activation during mouse oocyte meiotic maturation. J Biol Chem 292, 13784–13794.

Mihajlovic, A.I., and Bruce, A.W. (2017). The first cell-fate decision of mouse preimplantation embryo development: integrating cell position and polarity. Open Biol 7.

Mohammed, H., Hernando-Herraez, I., Savino, A., Scialdone, A., Macaulay, I., Mulas, C., Chandra, T., Voet, T., Dean, W., Nichols, J., et al. (2017). Single-Cell Landscape of Transcriptional Heterogeneity and Cell Fate Decisions during Mouse Early Gastrulation. Cell Rep 20, 1215–1228.

Moretto Zita, M., Soncin, F., Natale, D., Pizzo, D., and Parast, M. (2015). Gene Expression Profiling Reveals a Novel Regulatory Role for Sox21 Protein in Mouse Trophoblast Stem Cell Differentiation. J Biol Chem 290, 30152–30162.

Moriwaki, K., Tsukita, S., and Furuse, M. (2007). Tight junctions containing claudin 4 and 6 are essential for blastocyst formation in preimplantation mouse embryos. Dev Biol 312, 509–522.

Morris, S.A., Grewal, S., Barrios, F., Patankar, S.N., Strauss, B., Buttery, L., Alexander, M., Shakesheff, K.M., and Zernicka-Goetz, M. (2012). Dynamics of anterior-posterior axis formation in the developing mouse embryo. Nat Commun 3, 673.

Nichols, J., Silva, J., Roode, M., and Smith, A. (2009). Suppression of Erk signalling promotes ground state pluripotency in the mouse embryo. Development 136, 3215–3222.

Nichols, J., and Smith, A. (2009). Naive and primed pluripotent states. Cell Stem Cell 4, 487–492.

Nishioka, N., Inoue, K., Adachi, K., Kiyonari, H., Ota, M., Ralston, A., Yabuta, N., Hirahara, S., Stephenson, R.O., Ogonuki, N., et al. (2009). The Hippo signaling pathway components Lats and Yap pattern Tead4 activity to distinguish mouse trophectoderm from inner cell mass. Dev Cell 16, 398–410.

Nowotschin, S., Setty, M., Kuo, Y.Y., Liu, V., Garg, V., Sharma, R., Simon, C.S., Saiz, N., Gardner, R., Boutet, S.C., et al. (2019). The emergent landscape of the mouse gut endoderm at single-cell resolution. Nature 569, 361–367.

Perera, M., Nissen, S.B., Proks, M., Pozzi, S., Monteiro, R.S., Trusina, A., and Brickman, J.M. (2022). Transcriptional heterogeneity and cell cycle regulation as central determinants of Primitive Endoderm priming. Elife 11.

Pijuan-Sala, B., Griffiths, J.A., Guibentif, C., Hiscock, T.W., Jawaid, W., Calero-Nieto, F.J., Mulas, C., Ibarra-Soria, X., Tyser, R.C.V., Ho, D.L.L., et al. (2019). A single-cell molecular map of mouse gastrulation and early organogenesis. Nature 566, 490–495.

Posfai, E., Petropoulos, S., de Barros, F.R.O., Schell, J.P., Jurisica, I., Sandberg, R., Lanner, F., and Rossant, J. (2017). Position- and Hippo signaling-dependent plasticity during lineage segregation in the early mouse embryo. Elife 6.

Qiu, C., Cao, J., Martin, B.K., Li, T., Welsh, I.C., Srivatsan, S., Huang, X., Calderon, D., Noble, W.S., Disteche, C.M., et al. (2022). Systematic reconstruction of cellular trajectories across mouse embryogenesis. Nat Genet 54, 328–341.

Redo-Riveiro, A., Al-Mousawi, J., Linneberg-Agerholm, M., Proks, M., Perera, M., Salehin, N., and Brickman, J.M. (2024). Transcription factor co-expression mediates lineage priming for embryonic and extra-embryonic differentiation. Stem Cell Reports 19, 174–186.

Rivron, N.C., Frias-Aldeguer, J., Vrij, E.J., Boisset, J.C., Korving, J., Vivie, J., Truckenmuller, R.K., van Oudenaarden, A., van Blitterswijk, C.A., and Geijsen, N. (2018). Blastocyst-like structures generated solely from stem cells. Nature 557, 106–111.

Rossant, J. (2018). Genetic Control of Early Cell Lineages in the Mammalian Embryo. Annu Rev Genet 52, 185–201.

Saykali, B., Mathiah, N., Nahaboo, W., Racu, M.L., Hammou, L., Defrance, M., and Migeotte, I. (2019). Distinct mesoderm migration phenotypes in extra-embryonic and embryonic regions of the early mouse embryo. Elife 8.

Schrode, N., Saiz, N., Di Talia, S., and Hadjantonakis, A.K. (2014). GATA6 levels modulate primitive endoderm cell fate choice and timing in the mouse blastocyst. Dev Cell 29, 454–467.

Seong, J., Frias-Aldeguer, J., Holzmann, V., Kagawa, H., Sestini, G., Heidari Khoei, H., Scholte Op Reimer, Y., Kip, M., Pradhan, S.J., Verwegen, L., et al. (2022). Epiblast inducers capture mouse trophectoderm stem cells in vitro and pattern blastoids for implantation in utero. Cell Stem Cell 29, 1102–1118 e1108.

Silva, J., Barrandon, O., Nichols, J., Kawaguchi, J., Theunissen, T.W., and Smith, A. (2008). Promotion of reprogramming to ground state pluripotency by signal inhibition. PLoS Biol 6, e253.

Sousa, E.J., Stuart, H.T., Bates, L.E., Ghorbani, M., Nichols, J., Dietmann, S., and Silva, J.C.R. (2018). Exit from Naive Pluripotency Induces a Transient X Chromosome Inactivation-like State in Males. Cell Stem Cell 22, 919–928 e916.

Spangler, A., Su, E.Y., Craft, A.M., and Cahan, P. (2018). A single cell transcriptional portrait of embryoid body differentiation and comparison to progenitors of the developing embryo. Stem Cell Res 31, 201–215.

Spruce, T., Plass, M., Gohr, A., Ray, D., Martinez de Lagran, M., Rot, G., Novoa, A., Burguera, D., Permanyer, J., Miret, M., et al. (2022). The X-linked splicing regulator MBNL3 has been co-opted to restrict placental growth in eutherians. PLoS Biol 20, e3001615.

Stuart, H.T., Stirparo, G.G., Lohoff, T., Bates, L.E., Kinoshita, M., Lim, C.Y., Sousa, E.J., Maskalenka, K., Radzisheuskaya, A., Malcolm, A.A., et al. (2019). Distinct Molecular Trajectories Converge to Induce Naive Pluripotency. Cell Stem Cell 25, 388–406 e388.

Sutherland, A. (2003). Mechanisms of implantation in the mouse: differentiation and functional importance of trophoblast giant cell behavior. Dev Biol 258, 241–251.

Suzuki, D., Okura, K., Nagakura, S., and Ogawa, H. (2022a). CDX2 downregulation in mouse mural trophectoderm during peri-implantation is heteronomous, dependent on the YAP-TEAD pathway and controlled by estrogen-induced factors. Reprod Med Biol 21, e12446.

Suzuki, D., Sasaki, K., Kumamoto, S., Tanaka, K., and Ogawa, H. (2022b). Dynamic Changes of Gene Expression in Mouse Mural Trophectoderm Regulated by Cdx2 During Implantation. Front Cell Dev Biol 10, 945241.

Takahashi, M., Makino, S., Kikkawa, T., and Osumi, N. (2014). Preparation of rat serum suitable for mammalian whole embryo culture. J Vis Exp, e51969.

Takaoka, K., Yamamoto, M., and Hamada, H. (2011). Origin and role of distal visceral endoderm, a group of cells that determines anterior-posterior polarity of the mouse embryo. Nat Cell Biol 13, 743–752.

Tam, P.P., and Loebel, D.A. (2007). Gene function in mouse embryogenesis: get set for gastrulation. Nat Rev Genet 8, 368–381.

Tanaka, S., Kunath, T., Hadjantonakis, A.K., Nagy, A., and Rossant, J. (1998). Promotion of trophoblast stem cell proliferation by FGF4. Science 282, 2072–2075.

Tarazi, S., Aguilera-Castrejon, A., Joubran, C., Ghanem, N., Ashouokhi, S., Roncato, F., Wildschutz, E., Haddad, M., Oldak, B., Gomez-Cesar, E., et al. (2022). Post-gastrulation synthetic embryos generated ex utero from mouse naive ESCs. Cell 185, 3290–3306 e3225.

Thomas, P., and Beddington, R. (1996). Anterior primitive endoderm may be responsible for patterning the anterior neural plate in the mouse embryo. Curr Biol 6, 1487–1496.

Thowfeequ, S., Fiorentino, J., Hu, D., Solovey, M., Ruane, S., Whitehead, M., Zhou, F., Godwin, J., Mateo-Otero, Y., Vanhaesebroeck, B., et al. (2024). An integrated approach identifies the molecular underpinnings of murine anterior visceral endoderm migration. Dev Cell.

Verrier, L., Davidson, L., Gierlinski, M., Dady, A., and Storey, K.G. (2018). Neural differentiation, selection and transcriptomic profiling of human neuromesodermal progenitor-like cells in vitro. Development 145.

Wang, W., Yang, J., Liu, H., Lu, D., Chen, X., Zenonos, Z., Campos, L.S., Rad, R., Guo, G., Zhang, S., et al. (2011). Rapid and efficient reprogramming of somatic cells to induced pluripotent stem cells by retinoic acid receptor gamma and liver receptor homolog 1. Proc Natl Acad Sci U S A 108, 18283–18288.

Wood, H.B., and Episkopou, V. (1999). Comparative expression of the mouse Sox1, Sox2 and Sox3 genes from pre-gastrulation to early somite stages. Mech Dev 86, 197–201.

Xu, Y., Zhao, J., Ren, Y., Wang, X., Lyu, Y., Xie, B., Sun, Y., Yuan, X., Liu, H., Yang, W., et al. (2022). Derivation of totipotent-like stem cells with blastocyst-like structure forming potential. Cell Res 32, 513–529.

Yang, X., Chen, D., Sun, Q., Wang, Y., Xia, Y., Yang, J., Lin, C., Dang, X., Cen, Z., Liang, D., et al. (2023). A live-cell image-based machine learning strategy for reducing variability in PSC differentiation systems. Cell Discov 9, 53.

Ying, Q.L., Wray, J., Nichols, J., Batlle-Morera, L., Doble, B., Woodgett, J., Cohen, P., and Smith, A. (2008). The ground state of embryonic stem cell self-renewal. Nature 453, 519–523.

Zhao, T., Fu, Y., Zhu, J., Liu, Y., Zhang, Q., Yi, Z., Chen, S., Jiao, Z., Xu, X., Xu, J., et al. (2018). Single-Cell RNA-Seq Reveals Dynamic Early Embryonic-like Programs during Chemical Reprogramming. Cell Stem Cell 23, 31–45 e37.

