## Supplementary figures and images for "Chemically induced cell plasticity enables the generation of high-fidelity embryo model"

### Figure S1

#### Figure S1. Related to Figure 1

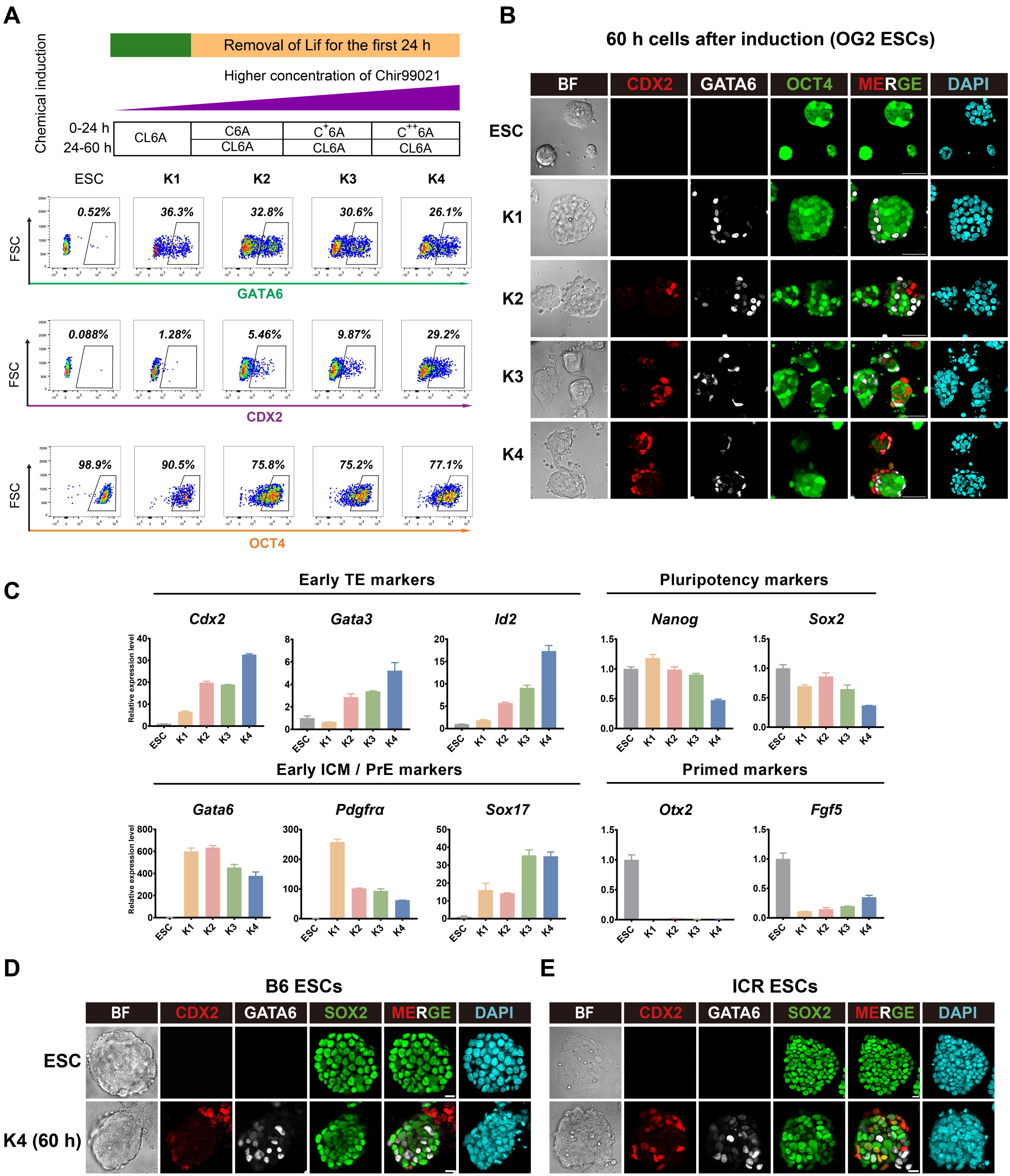

### Figure S2

#### Figure S2. Related to Figure 2

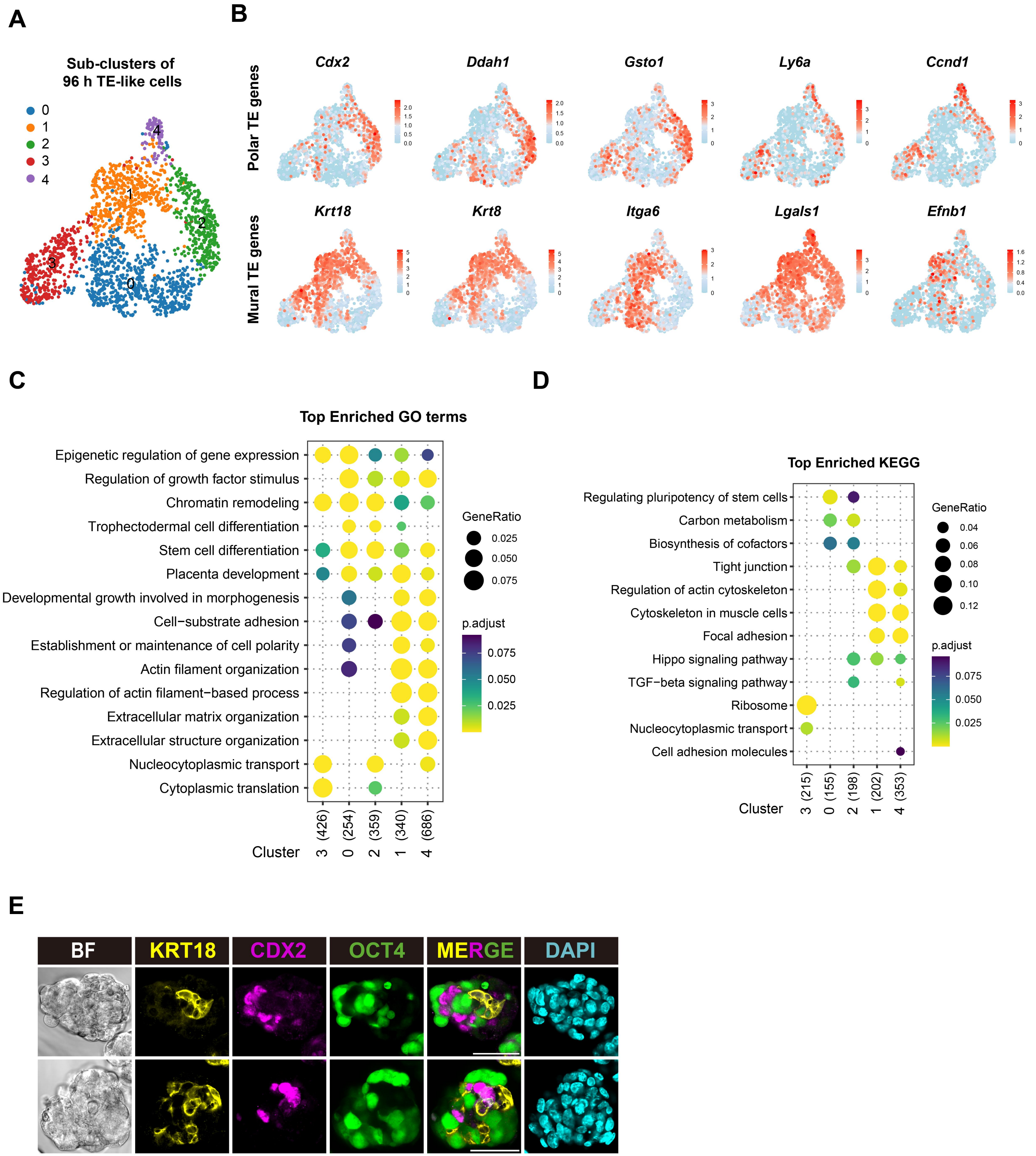

### Figure S3

#### Figure S3. Related to Figure 4

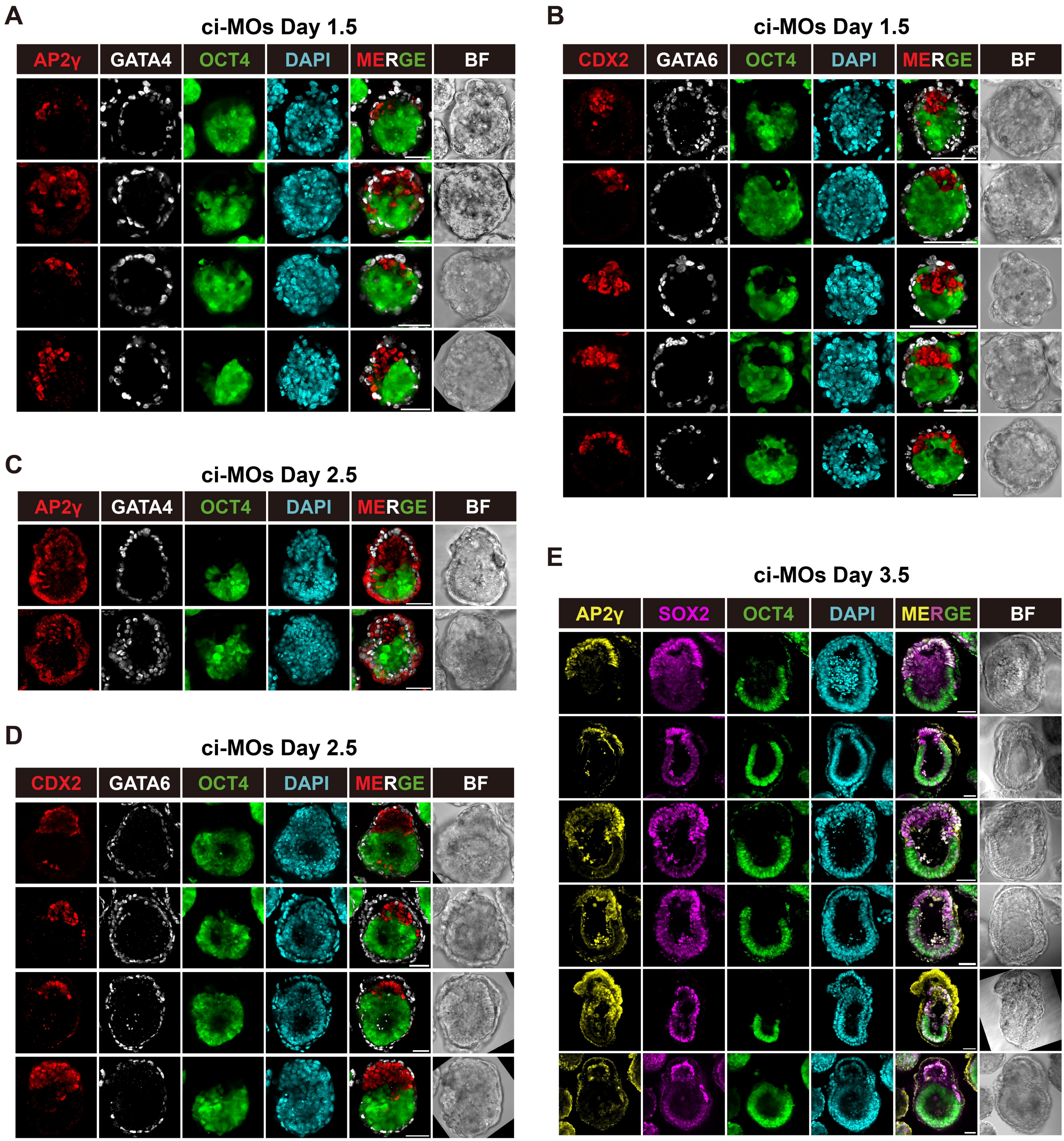

### Figure S4

#### Figure S4. Related to Figure 4

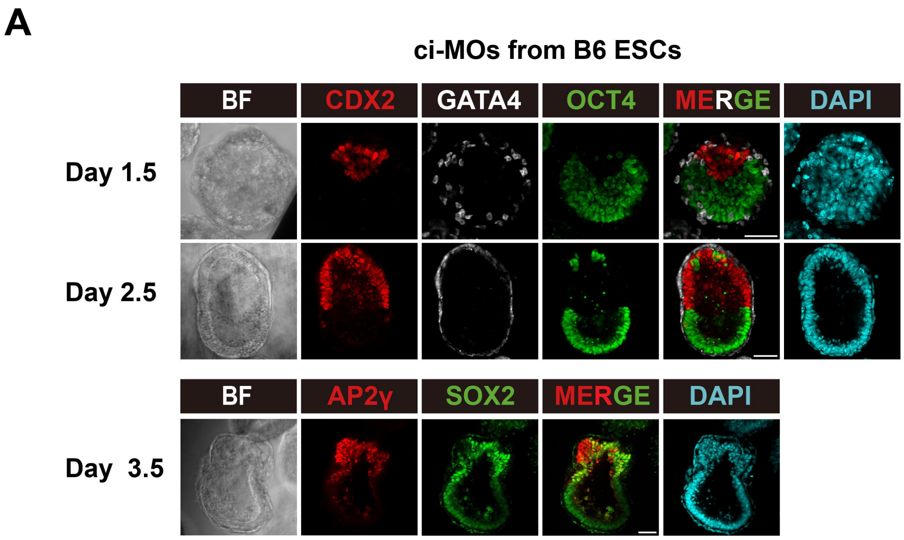


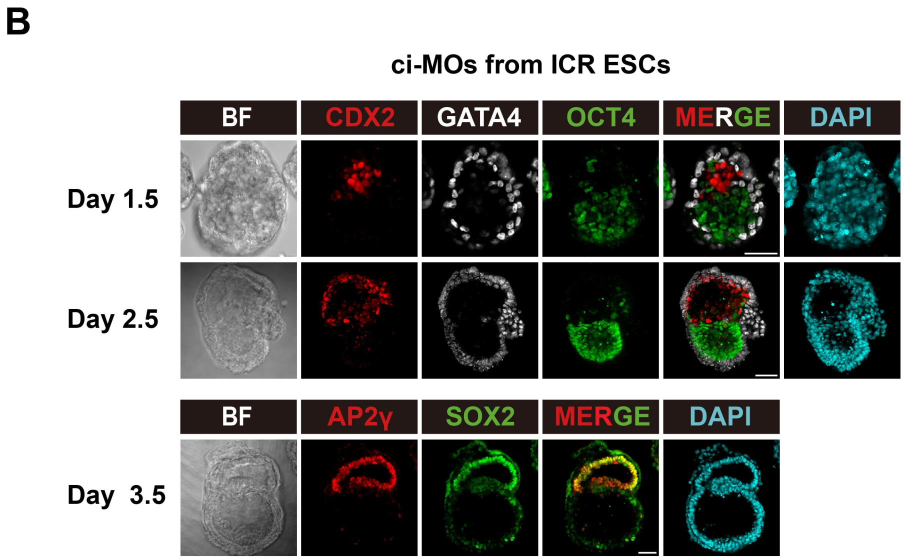

### Figure S5

#### Figure S5. Related to Figure 5

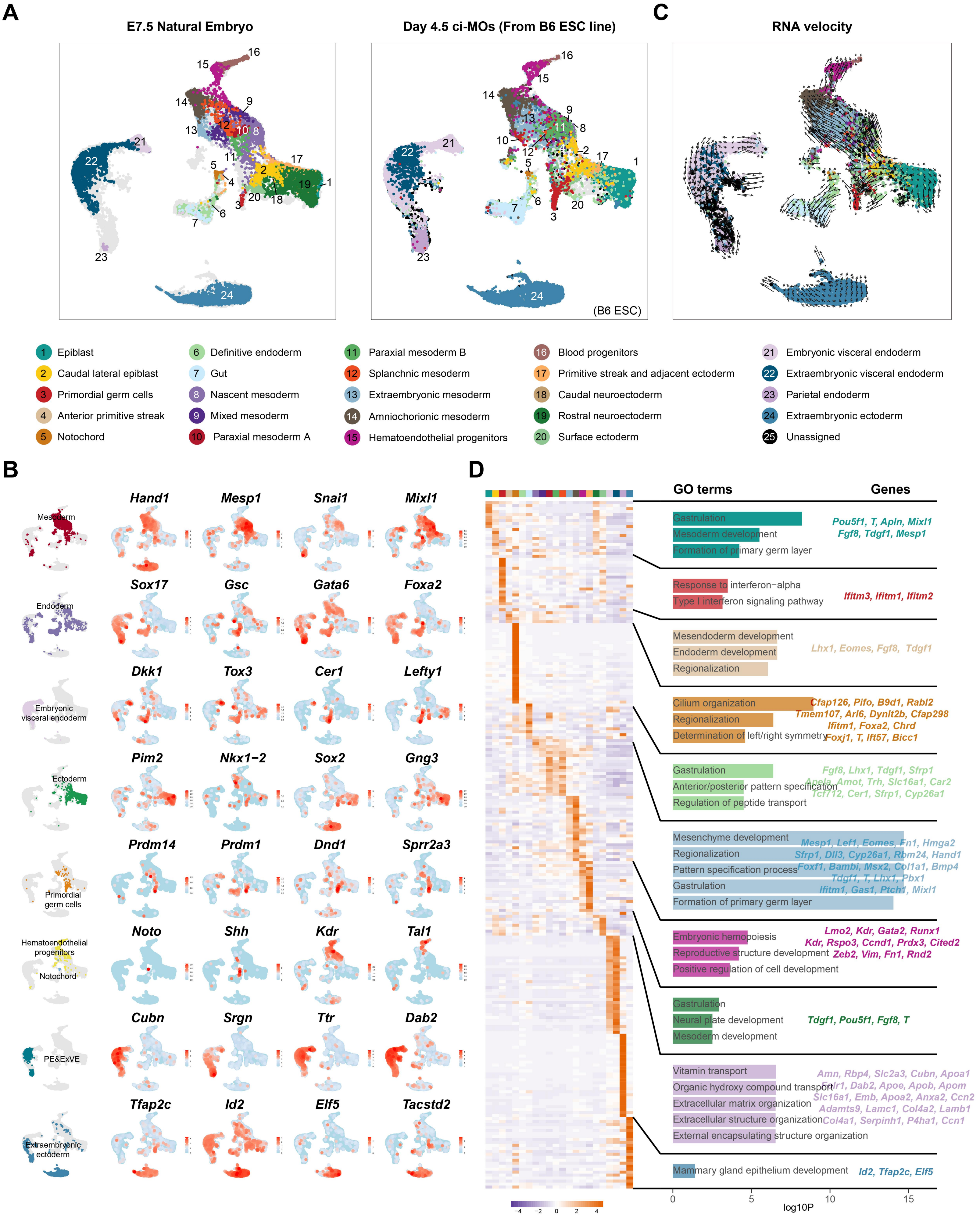
